# Epigenetic Profiling for the Molecular Classification of Metastatic Brain Tumors

**DOI:** 10.1101/268193

**Authors:** Javier I. J. Orozco, Theo A. Knijnenburg, Ayla O. Manughian-Peter, Matthew P. Salomon, Garni Barkhoudarian, John R. Jalas, James S. Wilmott, Parvinder Hothi, Xiaowen Wang, Yuki Takasumi, Michael E. Buckland, John F. Thompson, Georgina V. Long, Charles S. Cobbs, Ilya Shmulevich, Daniel F. Kelly, Richard A. Scolyer, Dave S. B. Hoon, Diego M. Marzese

## Abstract

Optimal treatment of brain metastases is often hindered by limitations in diagnostic capabilities. To meet these challenges, we generated genome-scale DNA methylomes of the three most frequent types of brain metastases: melanoma, breast, and lung cancers (n=96). Using supervised machine learning and integration of multiple DNA methylomes from normal, primary, and metastatic tumor specimens (n=1,860), we unraveled epigenetic signatures specific to each type of metastatic brain tumor and constructed a three-step DNA methylation-based classifier (BrainMETH) that categorizes brain metastases according to the tissue of origin and therapeutically-relevant subtypes. BrainMETH predictions were supported by routine histopathologic evaluation. We further characterized and validated the most predictive genomic regions in a large cohort of brain tumors (n=165) using quantitative methylation-specific PCR. Our study highlights the importance of brain tumor-defining epigenetic alterations, which can be utilized to further develop DNA methylation profiling as a critical tool in the histomolecular stratification of patients with brain metastases.

---

Brain metastases (BM) are the most common intracranial neoplasm in adults and are the next frontier for the management of metastatic cancer patients. Large population-based studies have shown that 8-10% of cancer patients develop brain metastases, with this proportion increasing up to 26% when autopsy studies were included^1-5^. Lung cancer, breast cancer, and cutaneous melanoma account for the vast majority (75-90%) of secondary neoplasms in the brain^1-4^. Treatment options for BM include surgery, whole-brain radiotherapy, stereotactic radiosurgery, and systemic-drug therapy, such as immunotherapy^6^. While systemic chemotherapy has limited efficacy, targeted therapies have recently shown promise for the management of patients with specific subtypes of cancer^6^. These tailored therapies have significantly affected treatment decision making for patients with breast cancer BM (BCBM). For example, patients with human epidermal growth factor receptor 2 (HER2)-positive BCBM can be treated with anti-HER2 agents^7^ and patients with estrogen receptor (ER)-positive BCBM can be treated with endocrine agents, cyclin dependent kinases 4 and 6 (CDK4/6) inhibitors, and the mechanistic target of rapamycin kinase (mTOR) inhibitors^8^. As such, accurate diagnosis is essential to effectively treat patients with metastatic brain tumors.

Diagnosis of BM is currently based on neuro-imaging and confirmed by anatomic pathology examination. When appropriate, the diagnostic algorithm begins by distinguishing BM from primary brain tumors using histologic features guided by the clinical and radiologic information^9^. Then, to identify the tissue of origin, morphological evaluation is supplemented by several immunohistochemistry (IHC) markers including thyroid transcription factor (TTF-1), chromogranin and synaptophysin for lung cancer BM (LCBM); GATA3 binding protein (GATA3), mammaglobin, gross cystic disease fluid protein 15 (GCDFP-15) and ER for BCBM; and human melanoma black 45 (HMB45), melanoma antigen recognized by T-cells 1 (Melan A/MART-1), SRY-Box 10 (SOX-10), and the S100 calcium binding proteins (S-100) for melanoma BM (MBM)^9,10^. However, a major limitation in achieving an accurate pathological diagnosis is the often poor differentiation and/or limited availability of metastatic brain tumor tissues to evaluate the complete panel of IHC markers^9^.

The advent of molecular classifiers based on the synergy between comprehensive tumor profiling and statistical modeling has dramatically improved the diagnosis, prognosis and, importantly, the therapeutic approaches for cancer patients^11^. To date, however, molecular classifiers to assist pathological diagnosis and improve stratification of patients with metastatic brain tumors have been ill-defined. DNA methylation (DNAm), an epigenetic modification of genomic DNA, was recently shown to be a powerful analytical tool to identify the origin of cancers from unknown primary sites^12^. We additionally have shown that DNAm profiling can be efficiently performed using small samples of BM tissues^13-16^. Here, we *hypothesized* that the construction and validation of DNAm classifiers for metastatic brain tumors could address current anatomic pathology diagnostic issues. The main objective of our study was to identify genomic regions whose DNAm status allow for *i)*- discrimination between primary and metastatic brain tumors, *ii)*- accurate identification of the tissue of origin for metastatic brain tumors, and *iii)*- can assist in the classification of therapeutically-relevant molecular subtypes for patients with metastatic brain tumors.

For this study, we included 165 patients with surgically resectable primary or metastatic brain tumors. Using Infinium HumanMethylation 450K (HM450K) microarray technology we generated high-quality genome-wide DNA methylomes for 96 microdissected BM specimens from patients with primary breast cancer (n=30), primary lung cancer (n=18), primary cutaneous melanoma (n=44), and patients with uncertain histogenesis of their brain tumor (n=4). We further integrated our data with additional publicly-available DNA methylomes (n=1,860) to construct and evaluate a robust brain metastasis DNAm classifier (**BrainMETH**). *BrainMETH* involves a three-step classification process that assists in accurately diagnosing brain neoplasms: first, by discriminating between primary and metastatic brain tumors (**Class A**), second, by identifying the tissue of origin of the BM (**Class B**), and finally, by providing information about molecular subtypes for patients with BCBM (**Class C**). We additionally used BrainMETH to identify the tumor site of origin of BM with an ‘uncertain’ diagnosis and to predict the status of IHC markers of BCBM that were not assessed in the initial diagnosis. Importantly, highlighting its potential clinical utility, we showed that targeted quantitative methylation-specific PCR (qMSP) can be applied to efficiently evaluate the representative genomic regions included in BrainMETH in DNA extracted from microdissected formalin-fixed paraffin-embedded (FFPE) archived tissue sections.

## Results

### Brain metastasis genome-wide DNA methylation data processing

To identify intrinsic differences in the epigenetic profiles of metastatic brain tumors, we first generated DNAm signatures for 96 microdissected BM tissues (a comprehensive list of BM specimens with detailed clinical and demographic information can be found as **Supplementary Data 1**). This comprehensive profiling was performed using the HM450K microarray, which, in addition to being the most commonly used genome-wide DNAm profiling platform currently, has also been employed by The Cancer Genome Atlas (TCGA) project to profile large cohorts of solid tumors^17,18^. Based on the most recent characterization of HM450K probes^19^, 102,941 genomic regions were excluded from downstream analyses. The exclusion criteria included probes that recognize common single nucleotide polymorphisms (SNPs), repetitive genomic elements, high GC density (>25%) on the 50-bp length probe sequence, a non-unique mapping to the genome, or low mapping quality (Fig. 1a). Additionally, to decrease potential biases associated with the gender of BM patients, we excluded 105,422 probes recognizing regions located in sex chromosomes or with proven cross-reactivity with sex chromosomes^20,21^. 2,983 probes with a detection P-value greater than 0.01 (‘NA’) in any specimen were excluded from this study. In addition, to obtain a set of genomic regions comparable to the newer generation of DNAm arrays, we excluded 13,882 probes that have been removed from the design of the HumanMethylation EPIC BeadChip array^22^. Finally, to decrease the influence of non-tumor cells, we excluded 48,797 genomic regions with no significant DNAm differences (Wilcoxon test; *P*-value >0.05) between BM specimens and normal brain tissues (n=100). Thus, this pipeline identified a dataset containing 211,552 informative HM450K probes to explore similarities and differences among brain tumors (**Fig. 1a**).

**Fig. 1:**
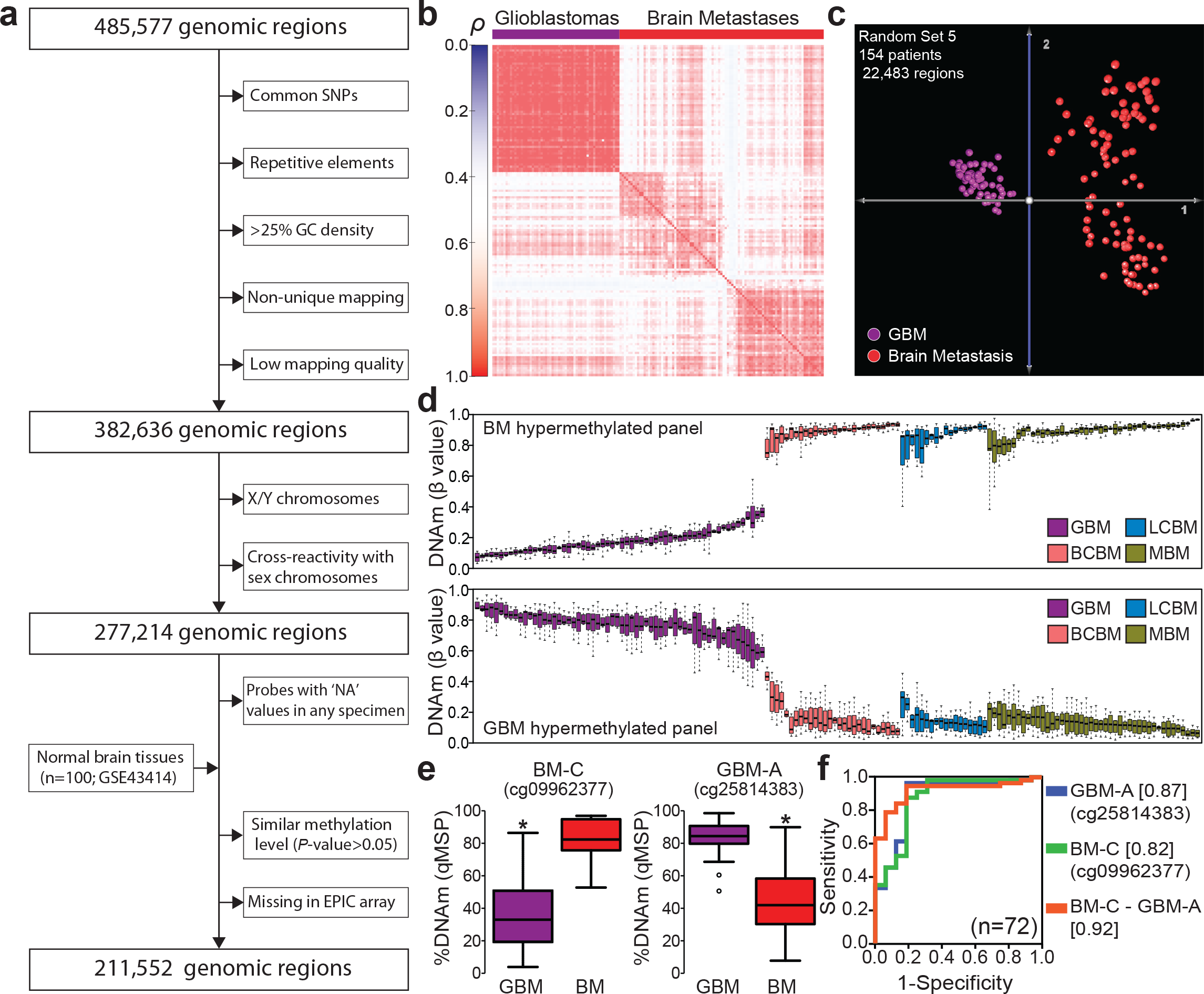
Genome-wide DNA methylation profiling of brain tumors. **a-** Diagram describing the normalization algorithm of Infinium HM450K probes. **b-** Matrix depicting the Spearman’s *ρ* correlation coefficients among primary and metastatic brain tumors. **c-** Principal component analysis (PCA) of GBM (n=60) and brain metastases (n=94) using DNA methylation level of 22,483 randomly-selected genomic regions. **d-** DNA methylation level of the 12 most differentially methylated genomic regions between GBM and BM specimens. The upper panel shows the β-value of six genomic regions differentially hypermethylated in BM (cg07076109, cg19111287, cg09962377, cg10982851, cg15002250, and cg23108580) and the lower panel shows the β-value of six genomic regions differentially hypermethylated in GBM (cg25814383, cg26306329, cg06663644, cg20604286, cg04314308, and cg26306994) for each specimen in the study (n=154). **e-** Validation of differentially methylated genomic regions by quantitative methylation-specific PCR (qMSP) in an independent cohort of brain tumor specimens (n=72). The left plot shows the methylation level of a genomic region hypermethylated in BM specimens (BM-C; cg09962377) and the right plot the level of a genomic region hypermethylated in GBM specimens (GBM-A; cg25814383; chr19:19,336,240). *Wilcoxon test, *P*-value <0.02. **f-** Receiver operating characteristic (ROC) curves showing the prediction potential of brain tumor type using qMSP scores for each independent genomic region and score for the combination (DNAm level of BM-C minus DNAm level of GBM-A; AUC =0.92, 95% CI=0.85-0.99); see **Supplementary Table 2** for details about validated genomic regions. Area under the curve values are indicated between square brackets.

### DNA methylation profiles of primary and metastatic brain tumors

Single intracranial metastases and primary brain tumors, mainly high-grade gliomas or glioblastomas (GBM), often exhibit overlapping clinical and radiological features^23^. To evaluate the potential utility of DNAm profiling for the classification of brain tumors, we identified a cohort of patients with GBM whose tumors have been profiled using the HM450K platform by TCGA-GBM project^24^. In order to reduce the impact of the lack of tissue microdissection on TCGA samples, we excluded GBM specimens with <70% tumor purity evaluated by the consensus purity estimation (CPE) method^25^. As expected, correlation and principal component analyses (PCA) using randomly selected HM450K probe sets (mean number of probes per set =22,543 ±150.7; **Supplementary Table 1**) revealed substantial DNAm differences between primary and metastatic brain tumors (**Fig. 1b-c**). The first three components of the PCA explained a mean of 84.14 ±0.6 percent of the cumulative variance (**Supplementary Table 1**). GBM specimens showed a more confined distribution in the three-dimensional component space than the BM specimens (**Fig. 1c**). Based on the intrinsic differences between DNAm distributions of primary and metastatic brain tumors, we explored differentially methylated genomic regions with the potential to discriminate between these tumor types. Employing a strict statistical cut-off (absolute DNAm difference >30%; False Discovery Rate (FDR)-corrected q-value <0.01), we identified 14,494 genomic regions differentially methylated between GBM and BM specimens (**Supplementary Fig. 1a**). Interestingly, the Genomic Regions Enrichment Annotations Tool (GREAT)^26^ indicated that regions hypomethylated in GBM (n=8,905) are predominantly associated with genes involved in neuronal differentiation and proliferation (Hypergeometric test; FDR-corrected q-value =1.96e-39) and regions hypomethylated in BM (n=5,589) are associated with genes involved in neuronal formation, development, and differentiation (Hypergeometric test; FDR-corrected q-value =1.14e-10; **Supplementary Fig. 1b**). Twelve differentially methylated genomic regions were selected based on the large differences in DNAm levels of these regions in GBM compared to any of the metastatic brain tumor types. Six regions were consistently hypermethylated in BM specimens (called herein BM-A to BM-F) and six regions were consistently hypermethylated in GBM specimens (called herein GBM-A to GBM-F; **Fig. 1d** and genomic coordinates and nearby genes described in **Supplementary Fig. 1c**). Using the HM450K data, we found that by surveying at least two genomic regions hypermethylated in BM we could distinguish primary from metastatic brain tumors with significant accuracy (AUC >0.90) (**Supplementary Fig. 1d**). Importantly, discrimination power was improved when combining genomic regions with distinct methylation patterns in GBM and in BM specimens. The evaluation of one BM hypermethylated region with one GBM hypermethylated region showed higher prediction (AUC =0.99, 95% CI =0.99-1.00) than evaluation of single genomic regions (**Supplementary Fig. 1e**). We, therefore, evaluated the classification potential of these genomic regions in independent test cohorts (n=227) of clinically annotated patients with GBM (GSE85539), patients with low-grade gliomas (LGGs) from the EORTC-26951 phase III clinical trial (GSE48461), and patients with MBM (GSE44661) included in our previous studies^14^. We found that assessment of single genomic regions provides substantial discrimination potential between BM and GBM (n=168; AUC range: 0.97 to 0.99; **Supplementary Fig. 2a**) and moderate discrimination efficiency between BM and LGG (n=75; AUC range: 0.67 to 0.89; **Supplementary Fig. 2b**). To further evaluate the pairing of BM hypermethylated regions and GBM hypermethylated regions, we then specifically assessed the DNAm level of these 12 genomic regions using qMSP in an independent cohort of microdissected brain tumor tissues (n=72). Initially, we assessed the DNAm level of the selected regions in microdissected GBM and BM clinical specimens and confirmed significant differences between GBM and BM specimens (Wilcoxon test; *P*-value <0.02; **Fig. 1e** and **Supplementary Fig. 2c**). Thus, we identified genomic regions with poor, moderate, and good qMSP performance (primer sequences and performance listed in **Supplementary Table 2**). We found that evaluating regions with moderate and good qMSP performance accurately distinguished metastatic brain tumors from GBM specimens (AUC =0.87, 95% CI=0.76-0.98). Importantly, a combination of two regions with good performance (BM score =DNAm level of BM-C minus DNAm level of GBM-A) showed improved prediction potential (AUC =0.92, 95% CI=0.85-0.99; **Fig. 1f** and **Supplementary Fig. 2d**). Based on these data, genomic regions whose DNAm level (as assessed by qMSP) exhibited good predictive accuracy, were selected as the first step of the DNAm-based brain tumor classifier (***BrainMETH* class A; Supplementary Table 2**).

### DNA methylation differences among brain metastases from melanoma, breast, and lung cancer

We observed that DNAm profiles of metastatic brain tumors present significantly lower overall Spearman’s correlation coefficients (mean *ρ* =0.77 ±0.02) as compared to primary brain tumors (mean *ρ*=0.90 ±0.06; *P*-value =2.2e-16; **Fig. 2a** and **Supplementary Fig. 3a**). Biologically, this epigenetic variability recapitulates the diversity of the tissue of the BM lesions. Based on these observations, we further explored differences among DNAm signatures of intracranial metastases. Interestingly, unsupervised cluster analysis of the top 5,000 most variable genomic regions precisely separated all the BM specimens into two main clusters (Bootstrap value =100%; **Fig. 2b** and **Supplementary Fig. 3b**). The influence of the epigenetic landscape of the tissue of origin was reflected in the organization of the hierarchical tree. Cluster A included all the BM from patients with epithelial primary tumors (breast and lung carcinomas) and cluster B contained all the BM from patients with neuroectodermal primary tumors (cutaneous melanoma; **Supplementary Fig. 3b**). Similar results were observed even when the number of the most variable genomic regions was increased from 5,000 to 100,000 (**Supplementary Fig. 3c-f**). By grouping all the BM specimens according to the pathologically confirmed primary tumor of origin (n=90), we found 31,818 genomic regions differentially methylated among the three BM types (one-way ANOVA; Bonferroni adjusted P-value <0.05; **Supplementary Table 3**). Using this set of genomic regions, PCA showed a clear separation of BM specimens according to the tumor of origin (**Fig. 2c**). We additionally evaluated BM specimens from four female patients treated for LCBM, but with ambiguous IHC profiling or with a history of both, primary lung cancer and primary breast cancer. In accordance with the pathological presumptive diagnosis, these four BM specimens with “uncertain” diagnosis showed overlapping with pathologically confirmed LCBM specimens (**Fig. 2d**). Moreover, we found that independent of the patient’s gender, this set of genomic regions accurately classified BM specimens according to the tumor of origin in a sub-cohort of female patients (n=58; **Fig. 2e**). These results reflect the efficiency and potential utility of DNAm profiling in accurately identifying the tissue of origin of intracranial metastases.

**Fig. 2:**
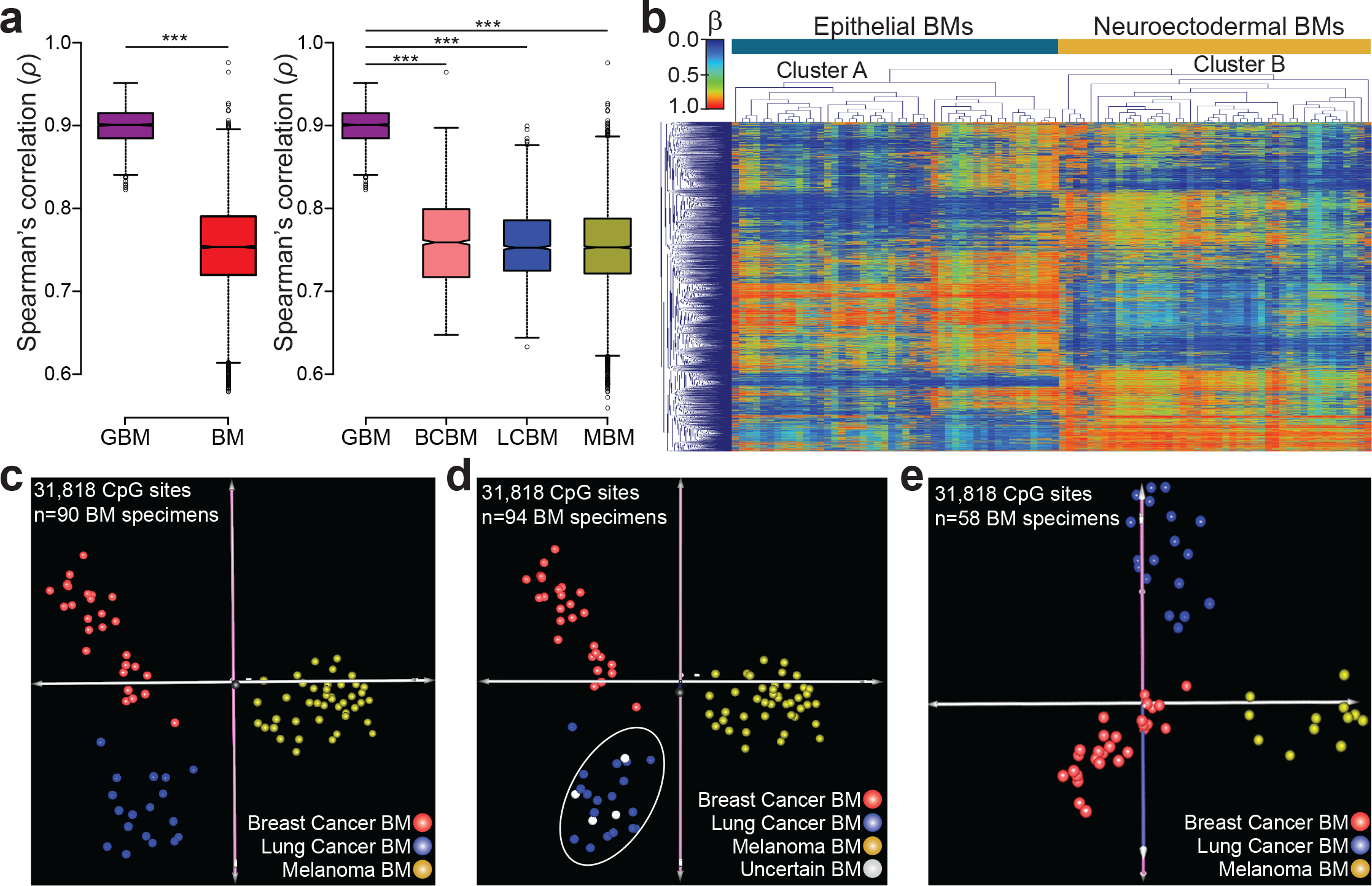
DNA methylation differences among brain metastases. **a-** Boxplot representing overall Spearman’s *ρ* correlation coefficients among brain tumors. The left plot includes all the GBM specimens (n=60, mean *ρ* =0.90 ±0.06) and all the BM specimens (n=94, mean *ρ* =0.77 ±0.02). The right plot describes the overall Spearman’s *ρ* correlation coefficient among BM specimens with anatomical pathology confirmed tumor of origin (BCBM n=28, LCBM n=18, and MBM n=44). \*\*\**P*-value <0.001. **b-** Unsupervised hierarchical clustering using Euclidean distance of the top 5,000 most variable genomic regions. **c-** Principal component analysis (PCA) using 31,818 CpG sites with significant (ANOVA, Bonferroni adjusted *P*-value <0.05; **Supplementary Table 3**) differential DNA methylation level among BM with anatomical pathology confirmed tissue of origin. **d-** PCA using the differentially methylated genomic region including four BM specimens with uncertain primary tumor of origin. **e-** PCA including BM specimens from female patients.

### DNA methylation classifiers to identify primary tumor origin of metastatic brain lesions

We constructed and evaluated DNAm classifiers to efficiently identify the tissue of origin of BM specimens. Our aim here was to find the minimum number of genomic regions capable of specifically predicting the origin of intracranial metastases. We employed a random forest (RF)-based supervised learning approach to construct classifiers for the tissue of origin of 96 BM specimens. We initially used the top 10,000 most variable differentially methylated genomic regions among BCBM, LCBM, and MBM specimens. Overall, the classifiers generated using this approach demonstrated an excellent classification potential (**Fig. 3a**) with an average sensitivity and specificity over 90% for all three BM types (melanoma, breast cancer, and lung cancer; **Fig. 3b**). We found that by surveying as few as 20 genomic regions, the classifiers exhibited median performance above 90%, with a deterioration of this value observed only when employing less than 10 regions (**Fig. 3a**). Thus, we identified the genomic regions with the highest importance for the prediction of the tumor of origin (**Fig. 3c**). These regions were tested in DNA methylomes from breast, lung and melanoma primary tumors generated by TCGA projects. We initially observed that the patterns of differential methylation of these regions partially agreed between our cohort of metastatic brain tumors and the TCGA cohorts (**Fig. 3d**). Interestingly, the top 100 most informative genomic regions showed good performance for the classification of primary tumor specimens according to the tumor of origin (**Supplementary Fig. 4a-b**). The first three components of the PCA explained 75.5% of the cumulative variance. These findings suggest that the regions with the potential to classify BM according to the tumor of origin represent epigenetic signatures specific to each tumor type. Nine genomic regions with high classification potential were then selected based on their low variance within each BM type, and large difference in mean DNAm among BM types (**Supplementary Fig. 4c-d**). Individually, DNAm levels of these genomic regions demonstrated good performance for the identification of specific BM tumor of origin (**Supplementary Fig. 5a**). We then evaluated the DNA methylation levels using qMSP in metastatic brain tumor clinical specimens (n=59) and categorized these regions according into good, moderate, and poor qMSP performance (**Supplementary Table 4**). Importantly, the qMSP evaluation of these genomic regions showed significant correlation with the HM450K assay (Spearman’s *ρ*; *P*-value <0.001; **Supplementary Fig. 5b**) and significant differential DNAm among the BM types (Wilcoxon test; *P*-value <0.001; **Fig. 3e**). These genomic regions showed excellent prediction performance to identify the tissue of origin (mean AUC =0.97), which was slightly improved by combining the results into tumor origin-specific methylation scores (mean AUC = 0.99; **Fig. 3f**). Of note, the qMSP evaluation of three genomic regions (MBM-B, LCBM-C, and BCBM-C) confirmed the origin of the four BM specimens with ‘uncertain’ diagnosis as LCBM, as forecasted by HM450K data (**Supplementary Fig. 5c**). Thus, genomic regions with good qMSP classification performance were selected as the second step of the brain tumor DNAm classifier (***BrainMETH* class B; Supplementary Table 4**).

**Fig. 3:**
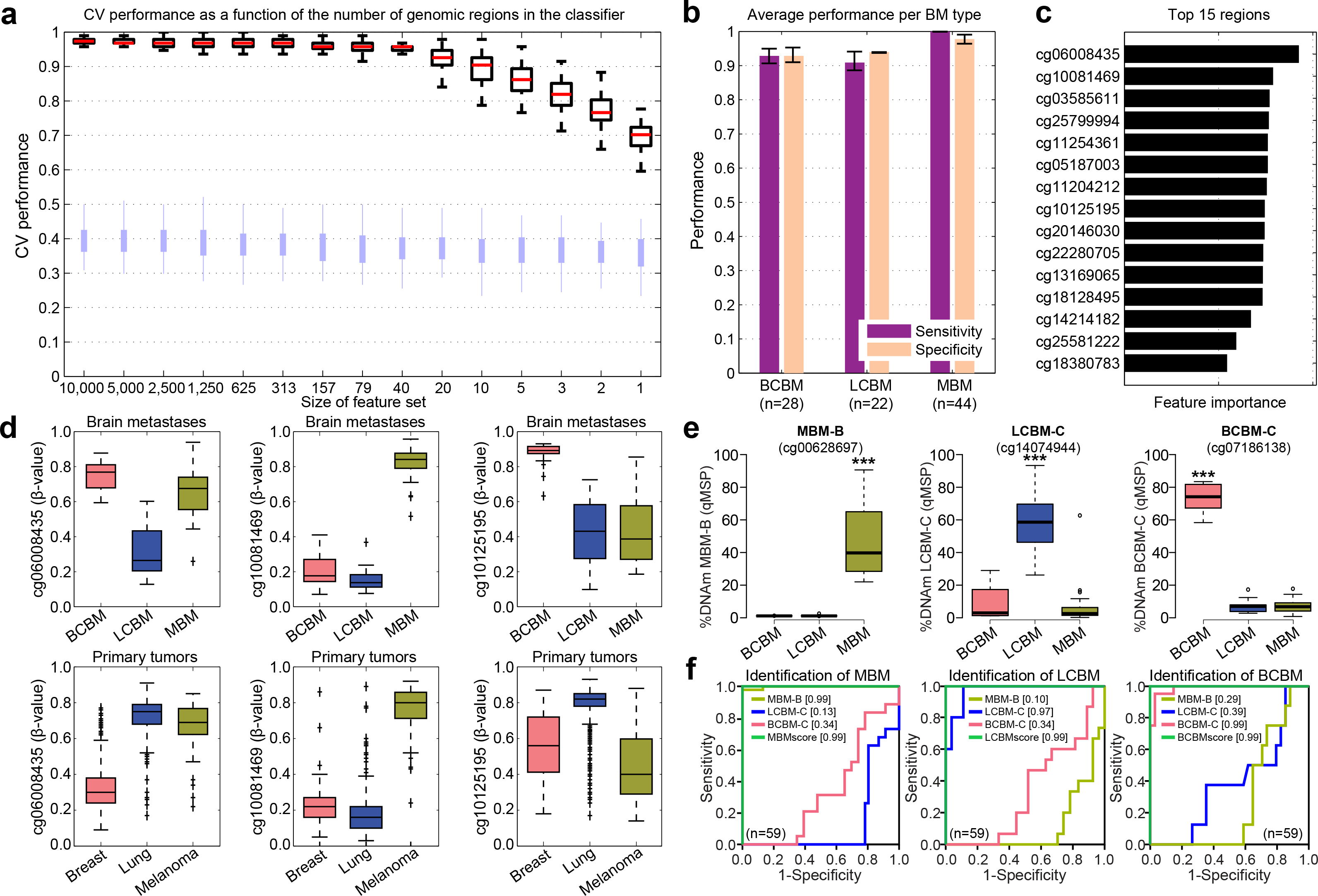
DNA methylation classifiers to predict the tissue of origin of metastatic brain tumors. **a-** Boxplots with cross-validation (CV) performance across 100 repeats for Random Forest (RF) classifiers to predict tumor of origin where, from left to right, decreasing numbers of features were used in model construction (x-axis). Red bars indicate medians and light blue bars depict the performance based on permuted class labels and represent the random background distribution. **b-** Bar plots depicting the prediction performance as measured by statistical sensitivity and specificity for each of the BM types. The bars show the average performance and interquartile range (error bars) across all models with 40 features or more across all repeats. **c-** Bar plots depicting the RF relative feature importance scores of the 15 most predictive genomic regions when summed across all models with 40 features or more and across all repeats. **d-** For three genomic regions in the top 15: boxplots of DNAm β-values across our cohort stratified by tumor of origin (BCBM n=28, LCBM n=22, and MBM n=44) in the upper panels and TCGA cohorts including primary breast cancer tumors (n=582), primary lung cancer tumors (n=379), and primary melanomas (n=83) in the lower panels. **e-** DNAm levels assessed by qMSP for three genomic regions differentially methylated among the three BM types. ***Wilcoxon test; *P*-value <0.001. **f-** ROC curves showing the prediction potential for the tumor of origin for each of the differentially methylated genomic regions and combinations into BM type-specific scores: **MBMscore =** DNAm level of MBM-B minus DNAm level of LCBM-C minus DNAm level of BCBM-C; **LCBMscore =** DNAm level of LCBM-C minus DNAm level of BCBM-C minus DNAm level of MBM-B; and **BCBMscore =** DNAm level of BCBM-C minus DNAm level of LCBM-C minus DNAm level of MBM-B; see **Supplementary Table 4** for details about these genomic regions. Area under the curve values are indicated between brackets.

### Breast cancer brain metastases molecular subtype-specific DNA methylation profiles

Characterization of the BM molecular subtype is critical to guide the clinical management of breast cancer patients. Therefore, we further explored the ability of DNAm profiling to identify molecular subtypes of BCBM specimens. The expression levels of the hormone receptors (HR; progesterone (PgR) and ER) and HER2 assessed by IHC, allowed us to classify 24 of the 28 BCBM into three clinically relevant classes: **1)** HR+/HER2-, **2)** HR-positive or HR-negative/HER2-positive (labeled HER2+), and **3)** HR-/HER2-. The first evaluation involved the identification of genomic regions differentially methylated among the three molecular subtypes. Hierarchical cluster analysis using 409 significantly differentially methylated regions (ANOVA; FDR-corrected q-value <0.0005; **Supplementary Table 5**) generated three sub-clusters containing each of the BCBM molecular subtypes (Bootstrap value =100%; **Fig. 4a**). Interestingly, HER2+ and HR+/HER2− BCBM specimens, which are generated from the late and differentiated luminal progenitor mammary cells, respectively, were included in a common cluster, while HR−/HER2− BCBM specimens, which commonly present gene expression profiles similar to mammary myoepithelial (basal layer) cells, were included in a separate cluster, indicating the relationship between the DNAm landscapes and the cell type of origin^27,28^, as well as supporting the concept that basal-like breast cancer represents a unique disease entity^29^. We found that these differentially methylated regions were organized in three specific clusters (**Fig. 4a** and **Supplementary Table 6**). The first cluster (**CL1**) included regions hypermethylated in HR+/HER2− BCBM and the third cluster (**CL3**) included regions hypermethylated in HR−/HER2− BCBM, while the second cluster (**CL2**) included regions hypermethylated in either HR+/HER2− or HER2+ BCBM and hypomethylated in HR−/HER2− BCBM specimens. This striking organization further highlights the relationship between DNAm profiles and the originating cell type (**Supplementary Table 6**). We then explored the associations between the three clusters of genomic regions and the specific BCBM molecular subtypes using a detrended correspondence analysis (DCA; cumulative inertia of the first three axes =80.1%; **Fig. 4b**). This analysis showed that not all these genomic regions are equally relevant to define the BCBM molecular subtypes. Therefore, by employing the nearest shrunken centroid algorithm, we identified 126 genomic regions specifically associated with BCBM molecular subtype (δ =2.1 and *ρ* =0.9; **Supplementary Table 7**). To gauge the ability of DNAm profiling to predict BCBM molecular subtypes, we included four additional patients without prior HR and/or HER2 IHC evaluation of their BM specimens, which were treated based on IHC evaluation of their primary breast cancer tumors and/or extra-cranial metastases. Multi-dimensional analysis using the DNAm level of the 126 genomic regions identified that two of these BCBM specimens (BCBM-10 and BCBM-23) belong to the HR+/HER2− subtype and the other two (BCBM-20 and BCBM-31) belong to the HER2+ subtype (**Fig. 4c**). Importantly, the molecular subtypes predicted by the DNAm analysis were further confirmed by the IHC analysis of ER, PR, and HER2 by the Clinical Laboratory Improvement Amendments (CLIA)-certified Department of Pathology at Providence Saint John’s Health Center (**Fig. 4d**). We then evaluated the utility of this epigenetic signature to recognize multiple BM lesions. To this end, we performed HM450K profiling of two cases with multiple BCBM. The first patient presented two synchronous brain metastatic tumors, and the second patient presented asynchronous metastatic lesions (BCBM-03 paired with BCBM-04 and BCBM-05 paired with BCBM-19, respectively; **Fig. 4e**). Interestingly, hierarchical analysis of DNAm profiles involving the 126 identified genomic regions (**Supplementary Table 7**) denoted an overlap between paired metastatic brain lesions. These results suggest that the BM molecular subtype can be inferred from the DNAm levels using a relatively small set of genomic regions.

**Fig. 4:**
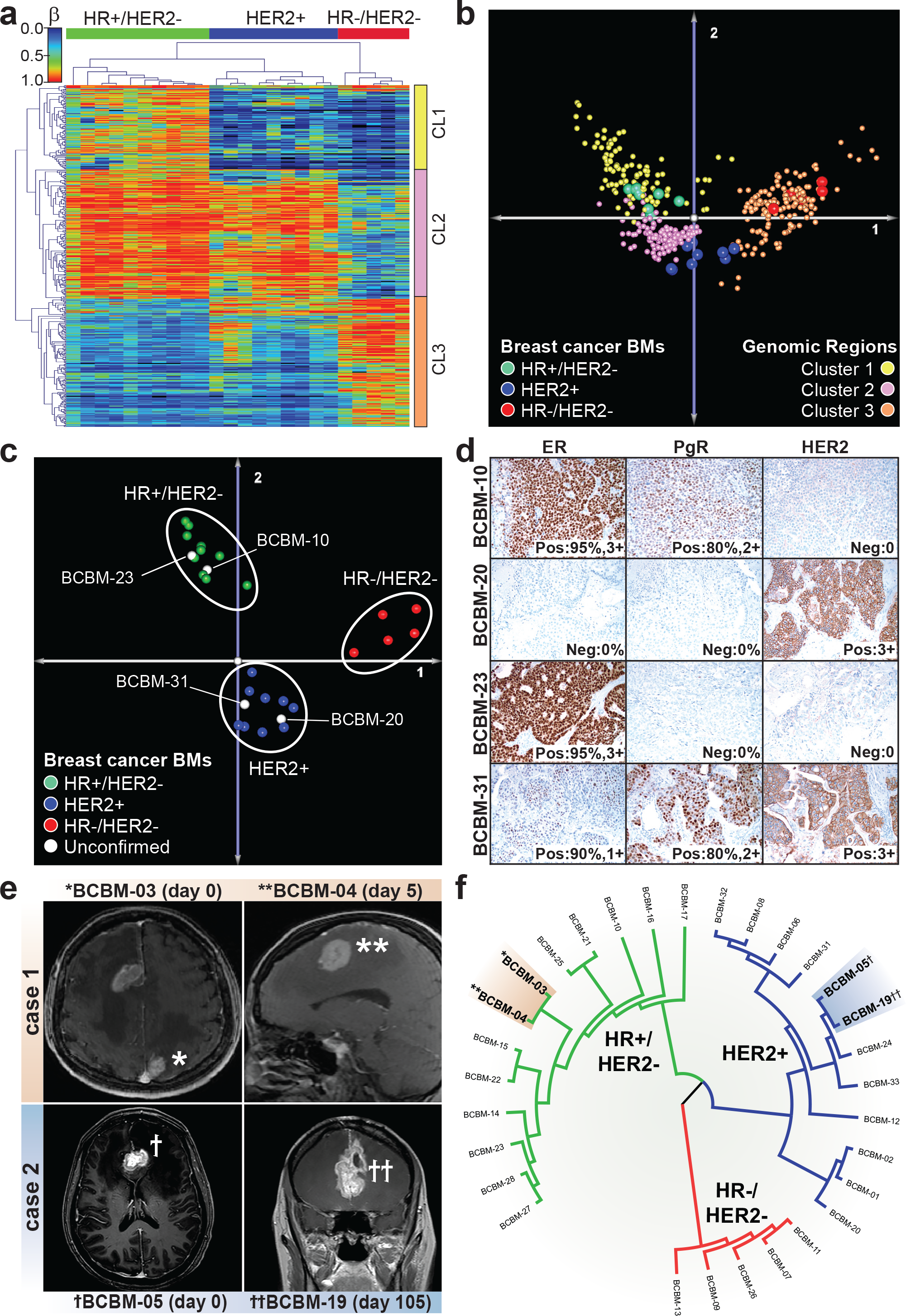
DNA methylation differences among breast cancer brain metastasis molecular subtypes. **a-** Hierarchical cluster analysis using Euclidean distance for the DNAm level of 409 genomic regions significantly differentially methylated (one-way ANOVA; FDR-corrected q-value <0.0005; Supplementary Table 5) among the three breast cancer molecular subtypes. This analysis revealed three distinct clusters of genomic regions. **CL1** includes regions specifically methylated in HR+/HER2− BCBM, **CL2** includes regions methylated in both, HR+/HER2− BCBM and HER2+ BCBM, and **CL3** includes regions specifically methylated in HR−/HER2− BCBM. **b-** Two-dimensional projection depicting the detrended correspondence analysis (DCA) for differentially methylated genomic regions (n=409) and BCBM specimens (n=24) with known IHC profile. This plot shows the spatial overlapping of genomic regions with relative importance for each BCBM molecular subtype. **c-** PCA including 24 BCBM specimens with known molecular subtypes and four BCBM with missing IHC information using 126 genomic regions with classification potential. The unconfirmed specimens were assigned to two different clusters. BCBM-10 and BCBM-23 overlapped with HR+/HER2− BCBM and BCBM-20 and BCBM-31 overlapped with HER2+ BCBM. **d-** IHC evaluation in a CLIA-certified Pathology Department for ER, PgR, and HER2 expression. The results confirm the DNAm-based prediction for the expression of HR and HER2. **e-** Magnetic resonance imaging showing two patients with synchronous (case 1) BCBM lesions (BCBM- and BCBM-04) and asynchronous (case 2) BCBM lesions (BCBM-05 and BCBM-19). **f-** Phylogenetic tree generated using the Euclidian metric distance for BCBM according to DNAm profile of the 126 genomic regions.

### Construction of DNA methylation classifiers to aid in the identification of breast cancer brain metastasis molecular subtypes

Due to the substantially different DNAm signatures observed among BCBM specimens, we set out to build an automated classification scheme that could accurately recognize therapeutically-orientated BCBM subtypes. Thus, our goals here were to study the prediction performance of DNAm-based classifiers and to assess whether classifiers with a small number of genomic regions are sufficiently powerful to discriminate between BCBM molecular subtypes. We employed an RF approach within a stratified cross-validation (CV) scheme to train and test a classifier that discriminates between the three molecular subtypes^30^. Starting with the top 10,000 most variable genomic regions, the classifiers demonstrated modest classification performance, showing improved performance compared to random classifications (**Fig. 5a-b**). Interestingly, the HR+/HER2− BCBM subtype could be predicted with the highest sensitivity and specificity (**Fig. 5b**). Moreover, classifiers using few probes retained good overall classification performance (**Fig. 5a**). Specifically, the median CV performance starts to deteriorate only with classifiers comprised of fewer than 10 genomic regions. Some of the probes with the highest feature importance scores were analyzed in detail through a comparison with the primary invasive breast cancer DNAm data generated by TCGA-BRCA project. The 15 highest ranked genomic regions all showed statistically significant differential methylation across the breast cancer molecular subtypes (**Fig. 5c**). The patterns of differential methylation of the most informative regions were in partial agreement between our study on BCBM and the primary breast cancer specimens cohort assessed by the TCGA-BRCA project (**Fig. 5d**). However, the genomic locations that efficiently classified BCBM according to specific molecular subtypes showed modest-to-poor classification efficiency for primary breast cancer specimens (**Supplementary Fig. 6a**). This result suggests that these genomic regions are exclusively relevant for molecular-subtype classification of BCBM specimens. Thus, we aimed to validate molecular subtype-specific DNAm signatures using qMSP. To this end, we again selected ten genomic regions with low variance within each molecular subtype, and large DNAm differences among the three classes (**Supplementary Fig. 6b-d**). These regions, alone or in combination, showed good discrimination potential for the respective BCBM molecular subtypes using the HM450K datasets (AUC >0.9; **Supplementary Fig. 7a**). We evaluated the DNAm level of these regions using qMSP in BCBM specimens (n=31). According to the qMSP performance each genomic region was classified into poor, moderate, and good performance (primer sequences and performance listed in **Supplementary Table 8**). We specifically focused on genomic regions with significant variation in the qMSP-based DNAm levels among the groups (Wilcoxon test; *P*-value <0.001; **Fig. 5e**) that exhibited a significant agreement with HM450K data (Spearman’s *ρ*; *P*-value <0.001; **Supplementary Fig. 7b**). Unlike the classification of BM according to the tumor of origin, which required one genomic region per class, we found that in order to accurately differentiate the BCBM molecular subtypes, at least two genomic regions per class were required (**Fig. 5f**). The combination of the DNAm levels of the final set of genomic regions presented a slightly higher predictive potential (mean AUC value =0.95) than single regions (mean AUC value =0.91) in all cases (**Fig. 5f**). Additionally, the qMSP analysis of these six genomic regions confirmed the BCBM molecular subtypes of the four specimens without IHC profile at the initial diagnosis (**Supplementary Fig. 7c**). Based on this evidence, this informative set of genomic regions validated by qMSP were selected for the DNAm-based classification of BCBM molecular subtypes (***BrainMETH* class C; Supplementary Table 8**).

**Fig. 5:**
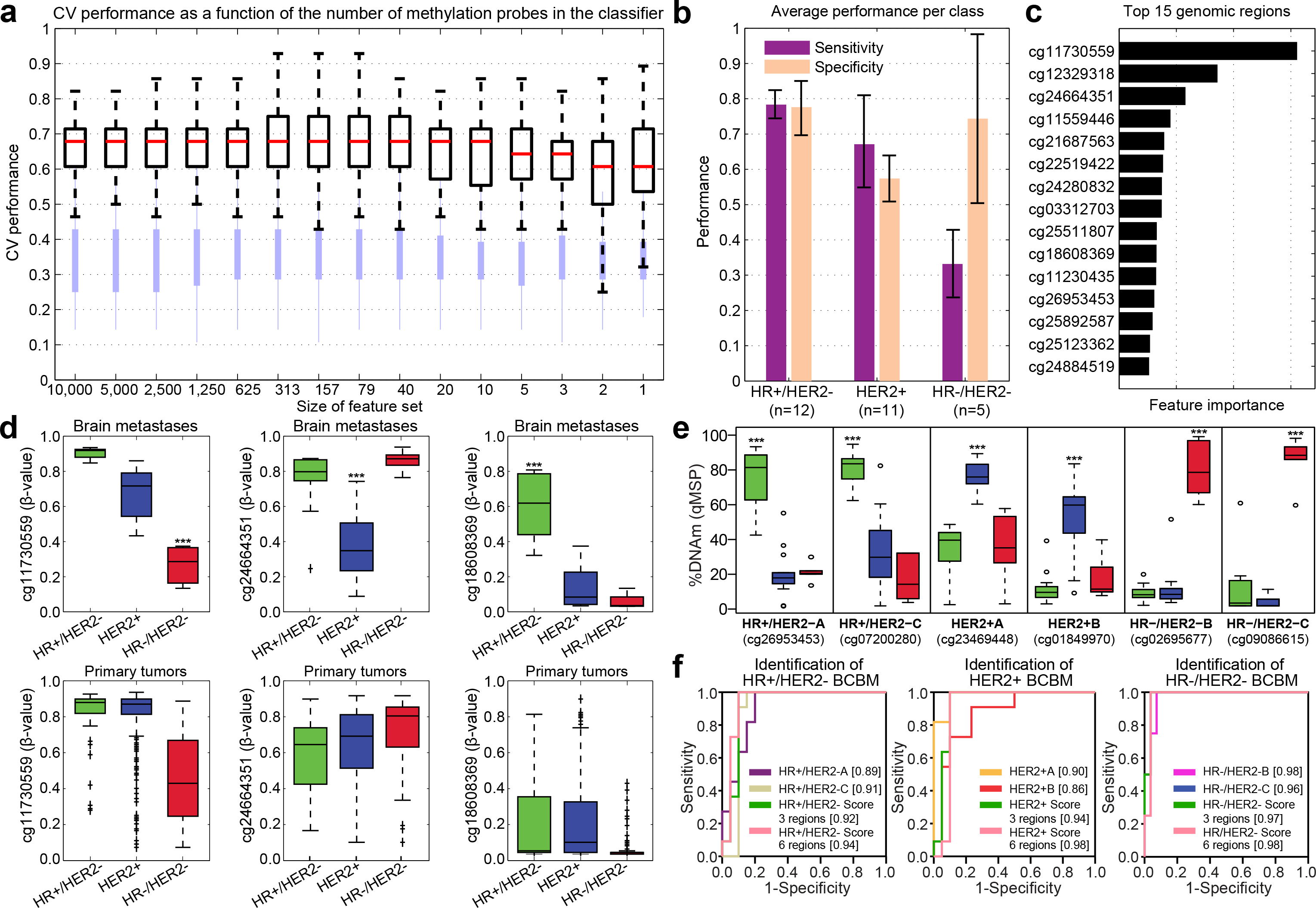
A DNA methylation-based classifier to predict molecular subtypes of breast cancer brain metastases. **a-** Boxplots with cross-validation (CV) performance across 100 repeats for RF classifiers to predict breast cancer molecular subtype. From left to right, decreasing numbers of features were used to construct the model (x-axis). Light blue bars depict the performance based on permuted class labels and represent the random background distribution. **b-** Bar plots depicting the prediction performance as measured by statistical sensitivity and specificity for each of the three molecular subtypes. The bars show the average performance and interquartile range (error bars) across all models with 10 features or more across all repeats. **c-** Bar plots depicting the RF relative feature importance scores of the 15 most predictive genomic regions when summed across all models with 10 or more features and across all repeats. **d-** Boxplots of DNAm levels (β-values) across our cohort stratified by molecular subtypes (HR+/HER2−; n=12, HER2+; n=11, and HR−/HER2−; n=5) and TCGA-BRCA cohort for primary breast cancer tumors stratified to match our molecular subtype definitions (HR+/HER2−; n=613, HER2+; n=137, and HR−/HER2−; n=164). **e-** DNAm levels assessed by qMSP for six genomic regions with differential DNAm among the three BCBM molecular subtypes. ***Wilcoxon test; *P*-value <0.001. **f-** ROC curves showing the prediction potential for the breast cancer molecular subtype for each of the six differentially methylated genomic regions and combinations of three or six genomic regions into BCBM molecular subtype-specific scores: **HR+/HER2−scores** = DNAm level of HR+/HER2− minus DNAm level of HER2+ minus DNAm level of HR−/HER2; **HER2+scores** = DNAm level of HER2+ minus DNAm level of HR+/HER2− minus DNAm level of HR−/HER2; and **HR−/HER2−scores** = DNAm level of HR−/HER2− minus DNAm level of HR+/HER2− minus DNAm level of HER2+; see **Supplementary Table 8** for details about these genomic regions. Area under the curve values are indicated between brackets.

## Discussion

In this study, we constructed and validated novel DNAm classifiers for the accurate diagnosis of brain metastases. Routinely, the first step in the diagnosis of metastatic brain tumors is to exclude a possible primary central nervous system neoplasm^9,31,32^. Here, we found substantial differences in the DNAm landscapes of primary and metastatic brain tumors. These findings are in concordance with a pilot study performed by Euskirchen et al. using nanopore sequencing^33^. Of note, our study found that the combination of small number of genomic regions provides high accuracy in discriminating primary from metastatic brain tumors. This is of great clinical and histopathological relevance, as we have demonstrated that these regions can be assessed by targeted qMSP using genomic DNA from routine FFPE clinical specimens. Additionally, this approach can be easily adapted to cover other genomic regions of interest to increase the robustness and expand the applications of the DNAm classifier. The genomic regions that we found useful in discriminating between GBM and BM have been included in the first step of the DNAm-based brain classifier (BrainMETH class A; summarized in **Fig. 6a**.

**Fig. 6:**
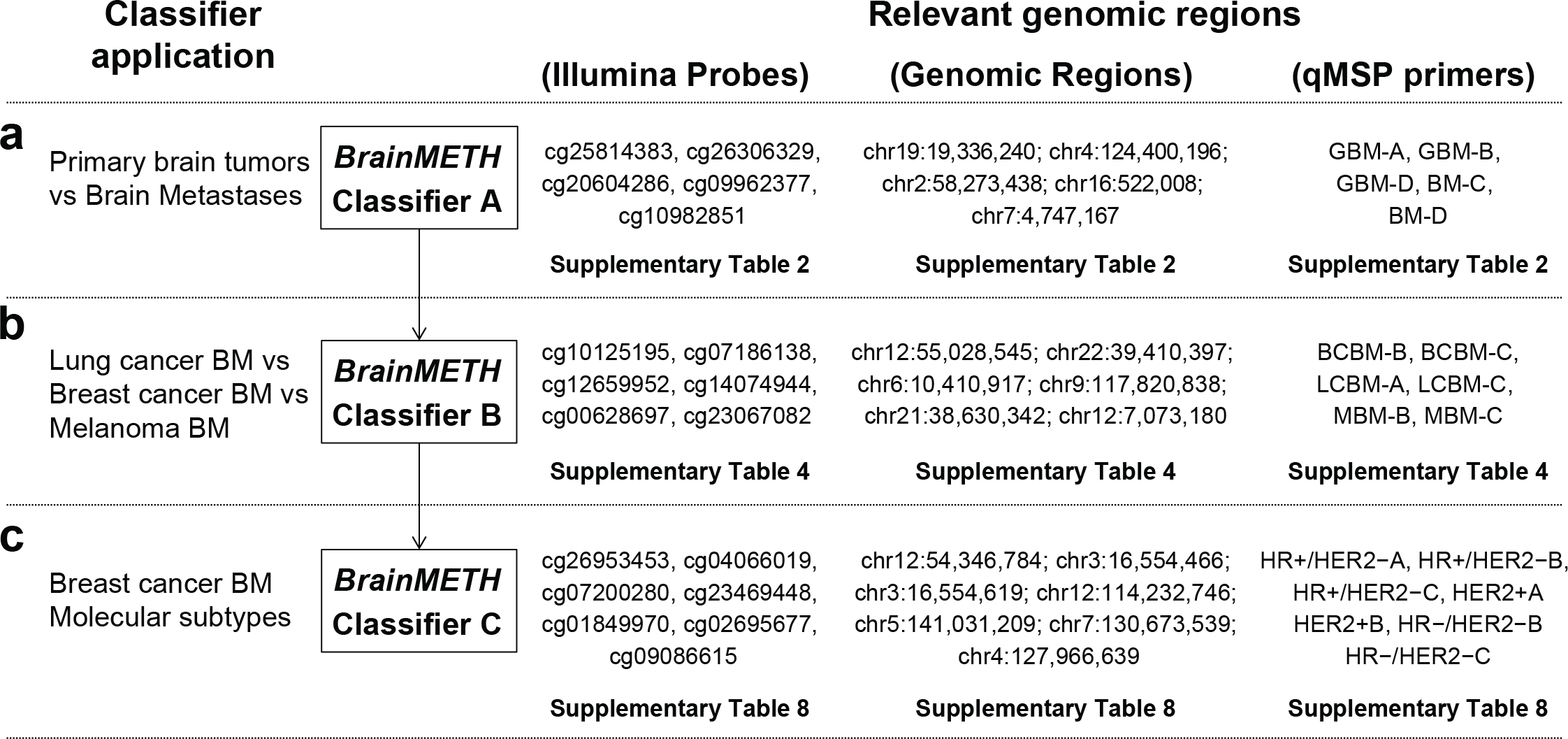
Summary of the BrainMETH classifiers. **a, b, c-** BrainMETH classifiers designed to discriminate between primary and metastatic brain tumors (**Classifier A**), among BM from different tumor of origin (**Classifier B**), and among BCBM from different molecular subtypes (**Classifier C**). A set of relevant Illumina probes, genomic regions, and validated primer sets by qMSP is provided for each step of the BrainMETH classifier.

In order to obtain an accurate prognosis and apply tumor-specific therapies, the second step in histopathology diagnosis for metastatic brain tumors is to identify the tumor of origin^6,34^. A recent study by Moran et. al. provided evidence about the utility of DNAm profiling to determine the origin of cancer of unknown primary^12^. In this regard, we found that BCBM, LCBM, and MBM present intrinsic differences in DNAm profiles that resemble the features of the embryonic origin for each cell type. Thus, MBM specimens, which are derived from the neuro-ectodermal layer, showed substantial differences compared to epithelial-derived BCBM and LCBM specimens. Using supervised learning by RF decision trees, we established a set of genomic regions that precisely classified BM according to the primary tumor of origin. In agreement with the presumptive histopathological diagnosis, BrainMETH accurately determined the origin of BM in patients with multiple primary tumors or with inconclusive anatomical pathologic evaluation. Interestingly, we found that these genomic regions also exhibited good classification performance of primary tumors from patients with melanoma, breast, and lung cancers. This observation suggests that CpG sites with good discrimination potential for BM overlap genomic regions associated with the tumor-type-specific epigenomic landscapes. Most importantly, we found that qMSP evaluation of three genomic regions using genomic DNA obtained from microdissected FFPE tissues showed excellent classification performance of BM specimens according to the tissue of origin. We therefore selected genomic regions with high discriminatory potential for the second step of the BM DNAm classifier (*BrainMETH* class B; **Fig. 6b**).

Finally, once the diagnosis of the tumor of origin has been established, it is crucial to identify molecular features that can stratify patients and guide therapeutic decisions. Undoubtedly, the expression of HR and HER2 are the most meaningful predictive factors for the treatment of breast cancer patients with metastatic brain tumors^35^. In clinical practice, BCBM therapeutically-relevant subtypes are usually inferred from the IHC profiling of the primary tumor or extra-cranial metastatic tissues. However, significant discrepancies have been recently demonstrated in the expression levels of ER and HER2 and the mutational landscape of primary and matched BCBM^36-38^. Therefore, if clinically feasible, it is currently recommended to perform a histopathological evaluation of the metastatic lesions to confirm the therapeutic subtype by at least reassessing the expression of HR and HER2^35^. To complement the conventional diagnosis, we explored the utility of DNAm profiling to classify BCBM into three therapeutically-relevant subtypes (HR+/HER2-, HER2+, and HR-/HER2-). Although the sample size of our BCBM cohort is relatively small, we were able to achieve good classification performance for all classes. We identified a set of genomic regions that accurately classified BCBM according to their pathologically confirmed molecular subtype. Remarkably, this same set of genomic regions successfully classified BCBM with initially unknown IHC profiling into clinically relevant subtypes. Most importantly, these classifiers only needed a small number of genomic regions to achieve good accuracy. As such, we conjecture that a qMSP assay for BCBM subtype determination would make a useful tool for clinical decision making. Thus, we found that the combination of as low as six genomic regions showed excellent classification potential for each BCBM subtype. To facilitate the development of the potential histomolecular application of the BrainMETH, we provide a list of genomic regions with good classification performance for BCBM therapeutic subtypes that can be inexpensively evaluated using FFPE tissues (*BrainMETH* class C; **Fig. 6c**).

In summary, our comprehensive identification and evaluation of the *BrainMETH* classifier emphasize that DNAm profiling has significant value in the molecular diagnosis of intracranial metastases. Indeed, data presented here demonstrate the potential of the *BrainMETH* classifier system to support conventional anatomic pathological evaluation. Yet, we have only examined BM originating from the three primary tumor types which most frequently metastasize to the brain. Further studies are required to assess the usefulness of DNAm classifiers in diagnosing patients with primary tumors with a less frequent incidence of BM, such as kidney, colorectal and ovarian cancers. Despite this need for a more expansive assessment, the findings of this study represent the first steps in developing a tool for accurate diagnosis, which is crucial to determining prognosis and guiding therapeutic decisions in patients with BM.

## Methods

### Patients and tissue specimens processing

In a multi-institutional effort, 165 patients with operable primary or metastatic brain tumors diagnosed at the Providence Saint John’s Health Center (Santa Monica, USA), Melanoma Institute of Australia (Sydney, Australia), and the Swedish Medical Center (Seattle, USA) were enrolled for this study. All clinical-demographic data and patient-derived samples were collected under research protocols approved by the joint Institutional Review Board of Providence Saint John’s Health Center/John Wayne Cancer Institute, the Western Institutional Review Board, the Swedish Institutional Review Board of Medical Center, and the Sydney Local Health District (Royal Prince Alfred Hospital Zone) Human Ethics Review Committee. All patients signed informed consent before joining the study. The experiments were performed in accordance with the World Medical Association Declaration of Helsinki and the National Institutes of Health Belmont Report. Tissues were de-identified and coded according to recommendations of the Health Insurance Portability and Accountability Act to ensure patient confidentiality.

### Histopathological evaluation and genomic DNA extraction

Representative FFPE tissue blocks for each BM lesion were selected by the respective Pathology Departments of the three institutions involved in the study. Neuropathologists reviewed tissue slides stained with hematoxylin & eosin for all specimens and identified areas with tumor cell enrichment (tumor purity) >70%. After deparaffinization, hematoxylin staining was performed in serial tissue sections. Eight-micrometer-thick tumor tissues were needle microdissected from consecutive FFPE slides. Genomic DNA was isolated using ZR FFPE DNA MiniPrep (D3066; Zymo Research, Irvine, CA), as previously described^13,14^.

### Genome-wide DNA methylation profiling and data access

Ninety-six BM specimens from 94 patients with BCBM (n=30), lung cancer (LCBM; n=18), cutaneous melanoma (MBM; n=44), and patients with both primary breast cancer and primary lung cancer (n=4), were evaluated using the Illumina Infinium HumanMethylation 450K BeadChips (HM450K; Illumina Inc., San Diego, CA, USA), as previously described^13,14^.

### Data availability

All HM450K raw and normalized data that support the findings of this study have been deposited in the NCBI’s Gene Expression Omnibus (GEO; https://www.ncbi.nlm.nih.gov/geo/) datasets under the series records GSE108576 (reviewer access token: upubgkywjxytjiv) and GSE44661.

### Quantitative-Methylation-Specific PCR

Sodium bisulfite modification was performed on 200-500 nanograms of genomic DNA using EZ DNA Methylation-Direct (D5021, Zymo Research, Irvine, CA, USA) following manufacturer recommendations. Target DNA methylation of GBM and BM genomic regions was performed using primers sets described in **Supplementary Tables S2, S4,** and **S6**. The quantitative amplification of methylated and unmethylated alleles was performed in CFX96 Touch™ Real-Time PCR detection system (185-5196; Bio-Rad Laboratories, Irvine, CA, USA), and the ΔCt (ΔCt = mean Ct methylated − mean Ct unmethylated) was calculated for each CpG site. The relative DNAm level was established by using the 2-ΔCt method, as previously described^14^. The percentage of DNAm was estimated by using a logarithmic equation derived from the analysis of a standard curve of serial dilutions of the universal methylated control (D5014, Zymo Research, Irvine, CA, USA) in universal unmethylated control (D5014, Zymo Research, Irvine, CA, USA) for each genomic region, as it has been shown^14^. Unless otherwise indicated in the text, BM type or subtype-specific DNAm scores were established as the average DNAm level for the BM type or subtype-specific region(s) minus the average DNAm level for the contrasting BM type or subtype-specific region(s).

### Histopathological and immunohistochemical evaluation of the BCBM

The BCBM specimens were evaluated at the CLIA-certified Department of Pathology, Providence Saint John’s Health Center, accredited by the College of American Pathologists (CAP). The BCBM samples were classified into molecular subtypes according to the expression levels of ER and PgR by IHC. HER2 was assessed by IHC and/or in situ hybridization assays. The FFPE tissue slides were sectioned at 4 μm and mounted on plus-coated glass slides, and immunohistochemically stained using a Ventana BenchMark ULTRA automated slide stainer (Roche Diagnostics, Indianapolis, IN, USA). Antibodies used were anti-ER (SP1, #790-4324, Ventana Medical Systems, Tucson, AZ, USA), anti-Progesterone Receptor (1E2, #790-2223, Ventana Medical Systems, Tucson, AZ, USA) and PATHWAY anti-HER-2/neu (4B5, #790-2991, Ventana Medical Systems, Tucson, AZ, USA). The scoring criteria considered for these markers were based on the ASCO/CAP guidelines^39,40^. Briefly, ER and PgR were considered positive if there was staining of the nucleus in at least ≥ 1% of the tumor cells in the sample. HER2 was considered positive for IHC 3+ or ISH amplified if single-probe average HER2 copy number >6.0 signals/cell or dual-probe HER2/CEP17 ratio ≥ 2.0. BCBM specimens were grouped according to the expression of these routinely clinically evaluated markers into a- HR+/HER2-, b- HR any/HER2+ (HER2+), and c- HR-/HER2-.

### Access to NCBI GEO DNA methylation datasets

HM450K data generated from 100 normal brain tissues; including 25 frontal cortex,
25 superior temporal gyrus, 25 entorhinal cortex, and 25 cerebellum specimens (GSE43414). Additionally, DNA methylomes generated from 152 GBM specimens (GSE85539) and 59 LGG specimens from patients enrolled in the phase III study EORTC 26951 (GSE48461). These data were integrated with the DNA methylomes from 16 melanoma BM that were previously published (GSE44661)^14,15^. These datasets were accessed using the R/Bioconductor ‘*GEOquery’* package v2.46.13, normalized using the SWAN method^41^ on the R/Bioconductor *‘wateRmelon’* package v1.19.1, and annotated using the R/Bioconductor *‘FDb.InfiniumMethylation.hg19’* package v2.2.0.

### Access to the Cancer Genome Atlas (TCGA) data and classification of patients

Clinical data for GBM, breast cancer, lung cancer, and cutaneous melanoma patients were downloaded from the Broad GDAC Firehose website (https://gdac.broadinstitute.org/) on April 2017. Genome-wide DNAm data for all these patients were retrieved from National Cancer Institute Genomic Data Commons Portal (https://gdc.cancer.gov/) using the R/Bioconductor *‘TCGAbiolinks’* package v1.2.5 ^42^.

### Selection of TCGA glioblastoma patients

We selected 60 patients with GBM from the TCGA-GBM project with an absolute tumor purity >70% and genome-wide DNAm level assessed by the HM450K beadchip platform. This cohort included patients with demographic characteristics paired with the cohort of patients with BM (**Table 1**) and a representative sampling of the GBM molecular subtypes.

### Selection of TCGA lung cancer and melanoma patients

We selected 380 primary lung cancer specimens corresponding to adenocarcinoma (n=152) and squamous (n=228) from the TCGA-LUAD and TCGA-LUSC projects, respectively. From the combined list of TCGA lung cancer patients (n=1,026), we excluded specimens obtained from patients with Stage IV (n=32), unknown stage (n=3), T4 from the AJCC 5^th^ or 6^th^ edition (n=27; since these cases can belong to the category of malignant pleural effusion currently considered M1a on the AJCC 7^th^ edition), recurrent sites (n=2), unavailable location (n=2), and tumor purity <70% (n=580). Additionally, we selected 83 primary cutaneous melanoma specimens from 466 cases included in the TCGA-SKCM project. Our exclusion criteria included specimens obtained from regional cutaneous or subcutaneous lesions including satellite and in-transit metastases (n=75), regional lymph node metastases (n=220), distant organ metastases (n=67), unavailable location (n=2), primary melanoma lesions from patients with stage IV (n=3), unavailable AJCC 7^th^ edition stage (n=2), and tumor purity <70% (n=16).

### Selection and classification of TCGA breast cancer patients

We identified and classified primary breast cancer specimens from the TCGA project with complete IHC information (n=914) and HM450K evaluation into 1-HR+/HER2- (n=613), 2-HRany/HER2+ (HER2+, n=137), and 3-HR-/HER2- (n=164). Briefly, from the 1,105 breast cancer samples included in the TCGA-BRCA database, we excluded specimens obtained from metastatic sites (n=7), male patients (n=12), unavailable gender information (n=2), and unavailable or indeterminate ER, PR or HER2 statuses (n=170). Cases with discordant results for HER2 expression by IHC and ISH were individually evaluated using the information provided in the pathological reports, and classified according to ASCO/CAP guidelines (n=15)^40^.

### Statistical and bioinformatics analyses

Tumor purity of all TCGA samples was assessed using the consensus purity estimation (CPE) method^25^. Differentially methylated genomic regions among the groups were identified using a one-way ANOVA. All *P*-values were two-sided and corrected for multiple comparisons using either Bonferroni or the FDR correction methods, as indicated in each case. DNAm data were dichotomized into ‘methylated’ (β-value ≥0.9) and ‘unmethylated’ (β-value ≤0.1) categories. Receiver operating curves (ROC) were used to estimate the sensitivity and specificity of the brain tumor classification method. The area under the curve (AUC) was calculated for each ROC to evaluate the accuracy of brain tumor classification based on DNAm observed in genomic regions. Hierarchical clustering (HCL) analyses were performed using Euclidean distance and average linkage clustering to identify relationships between genomic regions and brain tumors. Principal component analysis (PCA) was used to evaluate the overall variability, identify possibly correlated variables, and visualize the distance between selected brain tumor DNAm profiles. The robustness of phylogenetic tree reconstruction was determined by bootstrap resampling using 1,000 iterations using HCL analyzes. The HCL, PCA, terrain maps, and bootstraps analyzes were performed using the MultiExperiment Viewer v4.9^43^. Phylogenic trees were visualized using the FigTree Viewer v1.4.3. The multivariate detrended correspondence analysis (DCA) was used to find the most relevant genomic regions whose DNAm level was associated with each BCBM molecular subtype. DCA scores were established using the decorana function contained in the R *‘vegan’* package v2.3.5 and visualized using the COA function on the MultiExperiment Viewer v4.9^43^. The nearest shrunken centroids (NSCs) algorithm (initial parameters: delta >5, minimum correlation =0.5, number of bins =24) was applied to identify genomic regions that more accurately predict BCBM molecular subtypes. The NSCs were computed using the function pamr.train contained in the R *‘pamr’* package v1.55. DNAm classifiers to predict tumor of origin and BCBM molecular subtypes were trained and tested using the Random Forests (RF) algorithm^30^ applying the RF implementation for MATLAB v0.02 downloaded from http://code.google.com/pZrandomforest-matlab/. The RF classification models were run with 5000 trees each, default settings for the other parameters, and were trained and tested using a 3-fold stratified cross-validation (CV) strategy. Using only the samples in the training set, we first selected the 10,000 genomic regions which showed the largest variation in the median β-value across the different classes to train an RF model. Next, we selected the half of the genomic regions with the largest feature importance scores and retrained the RF using only this half of the regions. This process was iterated in 15 incremental steps, approximately halving the size of the feature set at each step, resulting in the 15th iteration in an RF classifier trained on only one genomic region. Each of the 15 RF classifiers was evaluated using the samples in the test set. This process was repeated using each of the three folds as the test set. Note that the test samples were never used for training. The whole procedure was replicated 100 times. The same procedure was repeated with permuted class labels for each of the 100 repeats in order to build a distribution of the performance of a ‘random’ classifier.

## Acknowledgements

We are grateful to Dr. Ian Hutchinson for his critical revision of the manuscript. This work was supported by the National Cancer Institute, National Institutes of Health (#R01CA167967 to D.S.B.H.; #P01CA077852 Core B to I.S. and T.A.K; and #U24CA210952 to T.A.K.); the Dr. Miriam and Sheldon G. Adelson Medical Research Foundation (to D.S.B.H.); the Ben and Catherine Ivy Foundation (P.H and C.S.C.); the AVON Foundation Breast Cancer Crusade (#02-2015-061 to D.S.B.H. and D.M.M.); the Associates for Breast and Prostate Cancer Studies (ABCs) award (#88737700140000 to J.I.J.O. and D.M.M.); and the Fashion Footwear Association of New York (FFANY) foundation award (#88737890550000 to J.I.J.O. and D.M.M.).

## Author contributions

J.I.J.O., T.A.K., D.S.B.H., and D.M.M conceptualized the study. J.R.J, Y.T., and M.E.B. performed the histopathological evaluation of tissue samples. G.B., J.F.T., G.V.L., C.S.C., and D.F.K. performed the selection of patients and clinical data annotation. J.S.W., P.H., and X.W. contributed with the tissue logistics and annotation of clinical data. D.M.M. designed the wet lab experiments. J.I.J.O. and A.O.M-P performed the wet lab experiments. T.A.K., M.P.S., I.S., and D.M.M performed the data normalization, statistical evaluations, and bioinformatics analyses. D.S.B.H. and D.M.M provided general guidance and oversaw the study. J.I.J.O., T.A.K., M.P.S., G.B., J.R.J, G.V.L., R.A.S., D.S.B.H., and D.M.M interpreted the results. J.I.J.O., T.A.K., A.O.M-P., and D.M.M. wrote the manuscript. All authors read and approved the final manuscript before submission.

## Supplementary Information

**Supplementary Figure 1:**
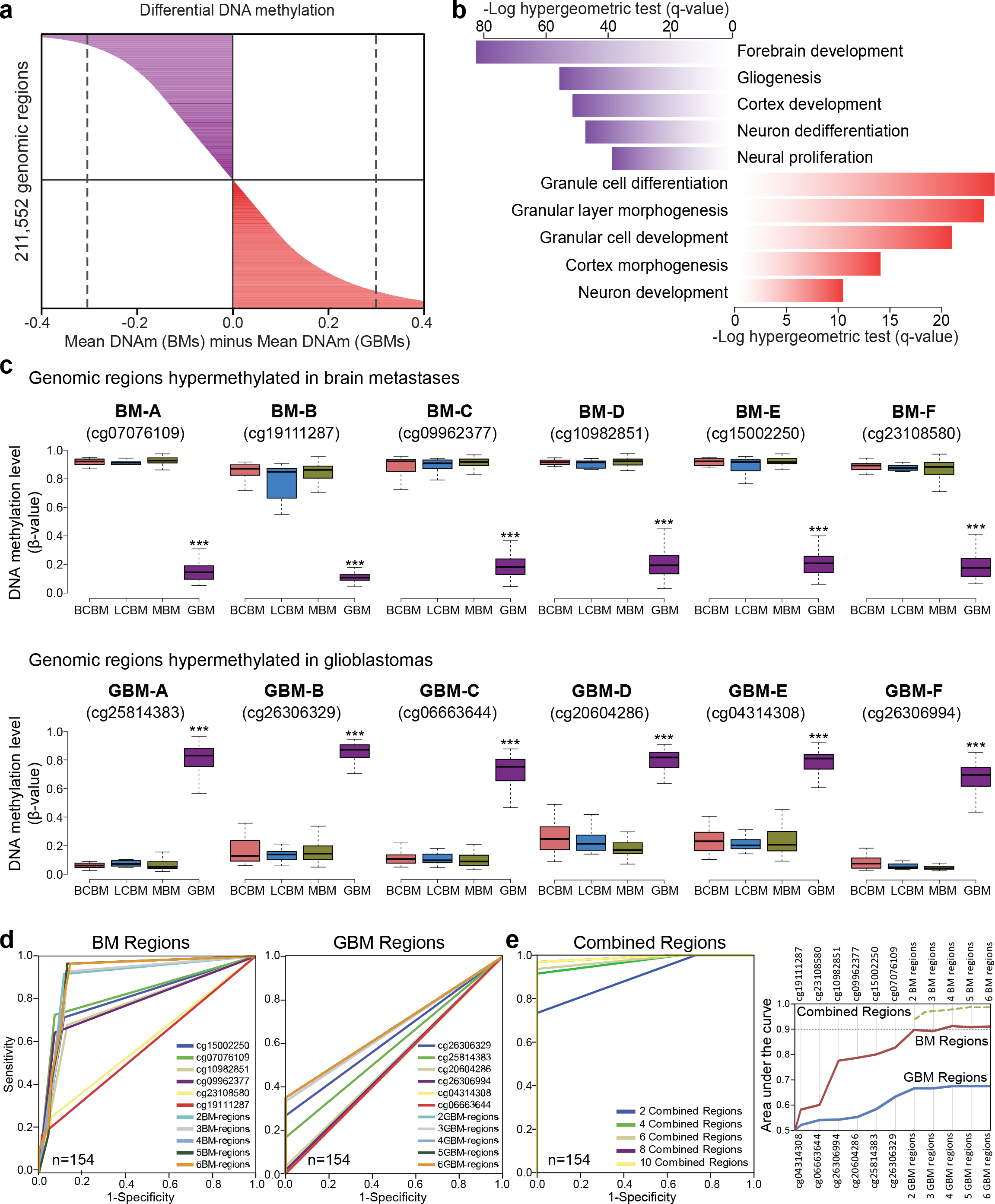
**a-** Differential DNA methylation between GBM and BM specimens. Mean β-values of 211,552 genomic regions in brain neoplasm specimens were compared between groups. Grey dashed lines demarcate genomic regions with at least 30% differential DNA methylation between the groups. **b-** Biological processes significantly enriched for genes with proximal (up to 5 kb) and distal (up to 500 kb) hypomethylated regions in GBM specimens (purple bars) and hypomethylated regions in BM specimens (red bars). **c-** Boxplots depicting the DNA methylation levels of the 12 genomic regions differentially methylated between primary and metastatic brain tumors (see **Supplementary Table 2** for details about the genomic location and distance to nearby genes). These regions were selected based on an overall low variance for DNA methylation level within each tumor type, and a large mean DNA methylation (HM450K microarray β-values) difference between primary (GBM, n=60) and metastatic (BCBM, n=28; LCBM, n=18; and MBM, n=44) brain tumors. The upper plots include genomic regions consistently hypermethylated in brain metastatic tumors (called herein BM-A to BM-F), and the lower plots include genomic regions consistently hypermethylated in primary brain tumors (called herein GBM-A to GBM-F). The differences in the DNA methylation levels between primary and metastatic brain tumors were considered statistically significant (Wilcoxon’s test) with a *P*-value less than or equal to 0.001 (***). **d-** Receiver operator curves (ROCs) showing the true positive rates (sensitivity) in function of the false positive rates (1-specificity) for the DNA methylation status of 12 genomic regions, six hypermethylated in metastatic brain tissues (**left plot**) and six hypermethylated in primary brain tumors (**right plot**), alone or in combination. In these analyses the DNA methylation levels (β-values) were dichotomized using a β-value cutoff into methylated (β-values ≥0.9) and unmethylated (β-values ≤0.1) statuses. **e-** ROCs for combinations of genomic regions distinctly hypermethylated in primary and metastatic brain tumors (**left panel**). Area under the curve (AUC) values (y-axis) for the individual and combined genomic regions specifically hypermethylated in primary brain tumors (GBM) and in metastatic brain tumors (BM; **right panel**).

**Supplementary Figure 2:**
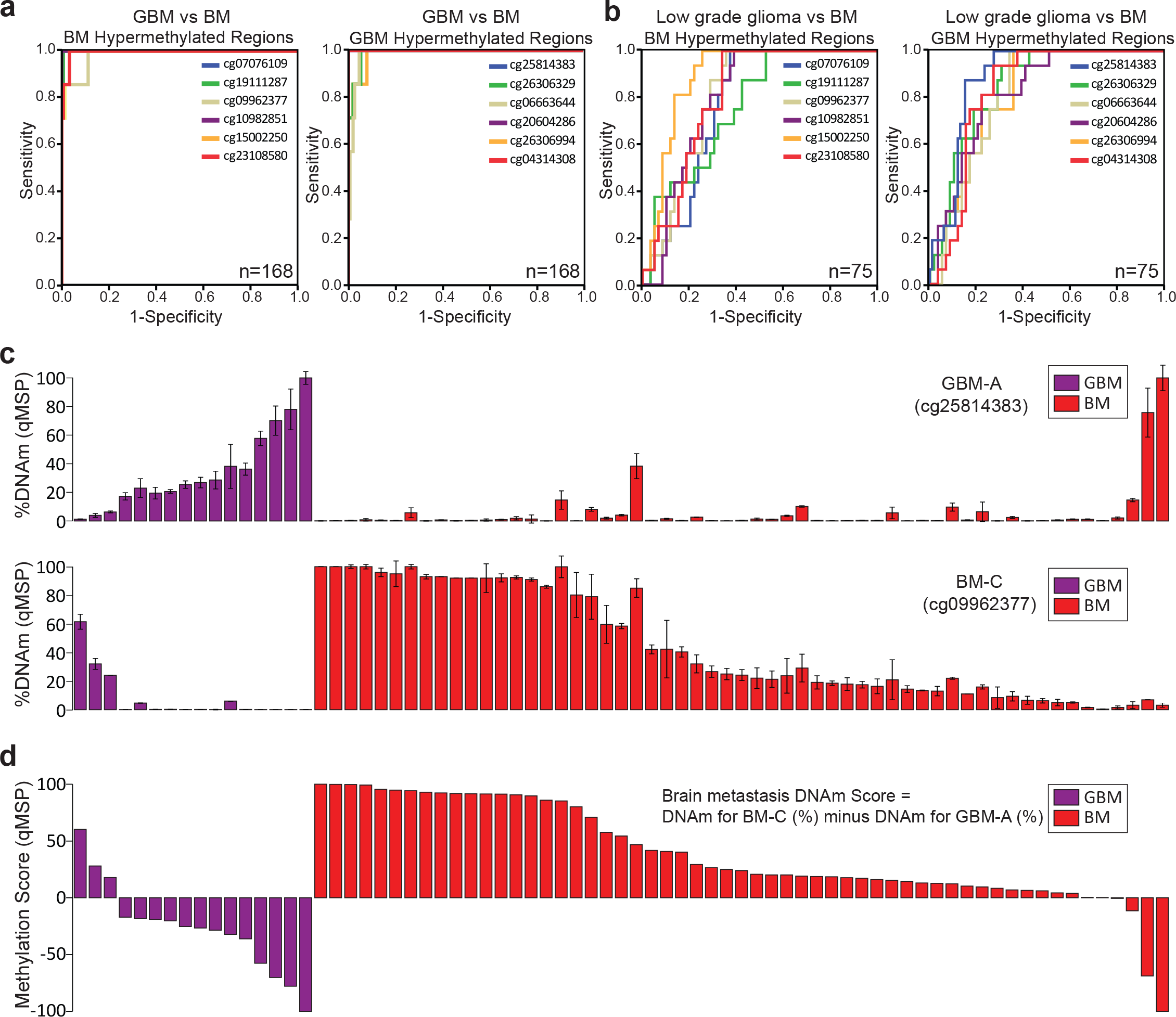
**a-** ROCs distinguishing between glioblastoma (GBM; GSE85539) and brain metastases (BM; GSE44661) for genomic regions hypermethylated in metastatic brain tumors (**left panel**) and genomic regions hypermethylated in GBM specimens (right panel). **b-** ROCs distinguishing between low-grade glioma (LGG; GSE48461) and BM (GSE44661) for genomic regions hypermethylated in metastatic brain tumors (**left panel**) and genomic regions hypermethylated in GBM specimens (**right panel**). **c-** Quantitative Methylation-Specific PCR (qMSP)-based DNA methylation level (%) for the most informative GBM-hypermethylated genomic region (GBM-A; **upper panel**) and the most informative BM-hypermethylated genomic region (BM-C; **lower panel**). Error bars represent the standard error of the mean (S.E.M.). **d-** Brain metastases DNA methylation score generated by subtracting DNA methylation level of a GBM genomic region (GBM-A) from the DNA methylation level of a BM genomic region (BM-C).

**Supplementary Figure 3:**
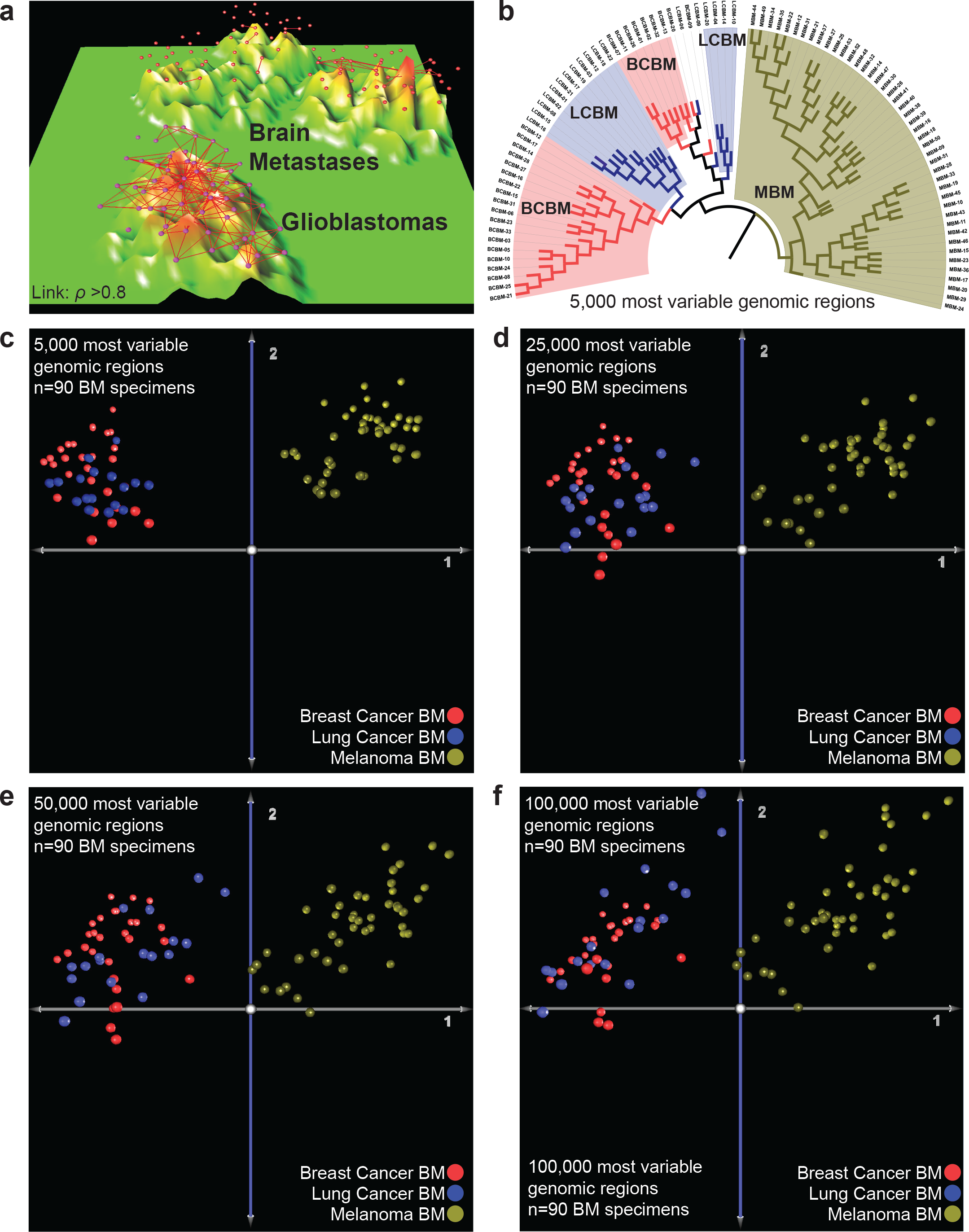
**a** Three-dimensional terrain map generated with the Spearman’s Rank Correlation coefficients (*ρ*) for all brain tumors presented in **Figure 2a**. Red spheres represent metastatic brain tumors (n=94), and purple spheres represent glioblastoma specimens (n=60). Red links indicate brain tumor pairs with a Spearman’s *ρ* correlation coefficient >0.8. **b-** Phylogenetic analysis of all the metastatic brain tumors with confirmed tissue of origin (n=90) using the DNA methylation levels of the top 5,000 most variable genomic regions. The phenetic tree was generated using Euclidean distances among all the metastatic brain tumor tissues. Major branches including brain metastases from the same origin were colored to visualize breast cancer (pink), lung cancer (light blue), and melanoma (brown). **c-f** Principal component analyses using the top 5,000 **(c)**, 25,000 **(d)**, 50,000 **(e)**, and 100,000 **(f)** most variable genomic regions for brain metastasis specimens with confirmed tumor of origin.

**Supplementary Figure 4:**
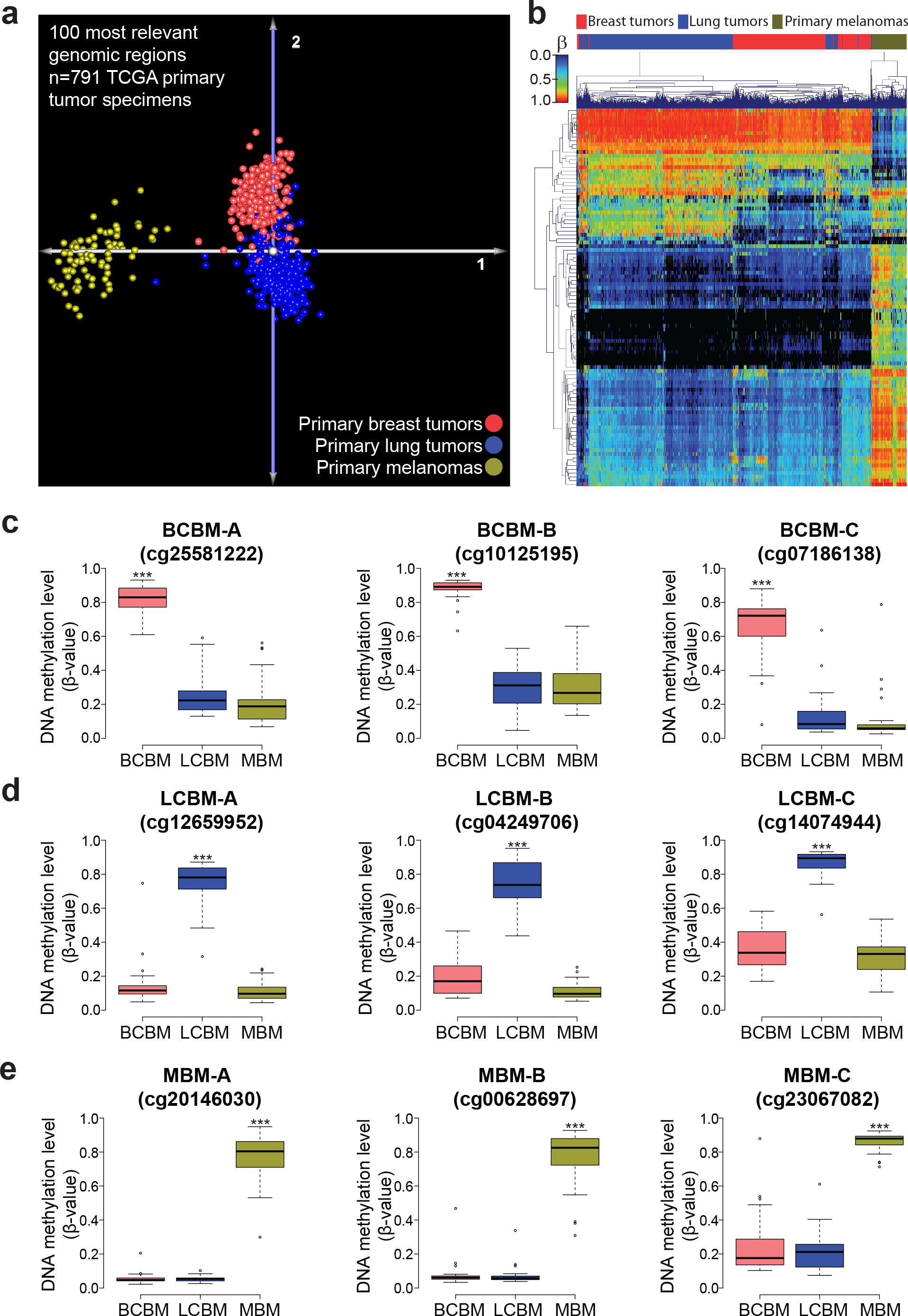
**a-** Principal component analysis for primary tumors from the breast (pink spheres; n=401), lung (blue spheres, n=307), and melanoma (brown spheres; n=83) using the top 100 most informative genomic regions for the classification of metastatic brain tumors. Only TCGA primary tumor specimens detection P-value greater than 0.01 for the 100 selected CpG sites were included in this analysis. **b-** Unsupervised hierarchical cluster analysis for TCGA primary tumor specimens using the top 100 most informative for the classification of metastatic brain tumors. **c-e** Boxplots showing the DNAm levels (HM450K microarray β-values) of nine genomic regions differentially methylated among the three types of brain metastases (see **Supplementary Table 4** for details about the genomic location and distance to nearby genes). This set of regions includes three CpG sites hypermethylated in breast cancer brain metastases (BCBM; **c**) specimens, three CpG sites hypermethylated in lung cancer brain metastases (LCBM; **d**), and three CpG sites hypermethylated in melanoma brain metastases (MBM; **e**). *** indicates a Wilcoxon’s test-based *P*-value less than or equal to 0.001.

**Supplementary Figure 5:**
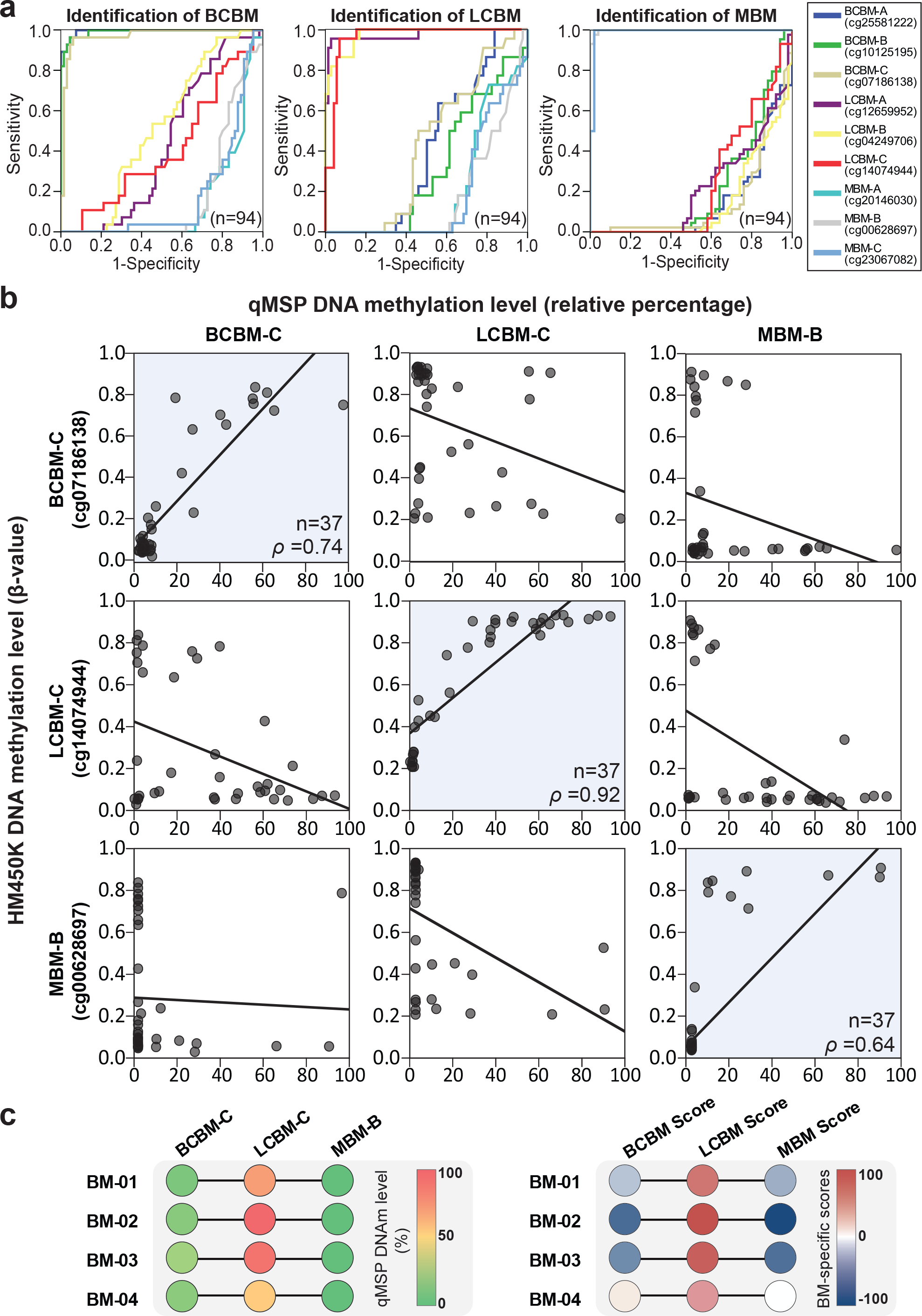
**a-** ROCs distinguishing between the three types of brain metastases using the DNA methylation levels (HM450K microarray β values) of the nine differentially methylated genomic regions (**Supplementary Table 4**). **b-** Spearman´s *ρ* correlation among DNA methylation levels assessed by HM450K (β-values, **y-axes**) and targeted approach (qMSP; **x-axes**) for the selected three genomic regions. **c-** qMSP analysis of the three selected genomic regions of the four samples with an uncertain primary origin of BM. The left panel shows the DNAm level (percentage) for each region. The right panel shows the DNAm scores specific for each type of BM. Each score was calculated as follow: **BCBMscore** = DNAm level of BCBM-C minus DNAm level of LCBM-C minus DNAm level of MBM-B; **LCBMscore** = DNAm level of LCBM-C minus DNAm level of BCBM-C minus DNAm level of MBM-B; and **MBMscore** = DNAm level of MBM-B minus DNAm level of LCBM-C minus DNAm level of BCBM-C.

**Supplementary Figure 6:**
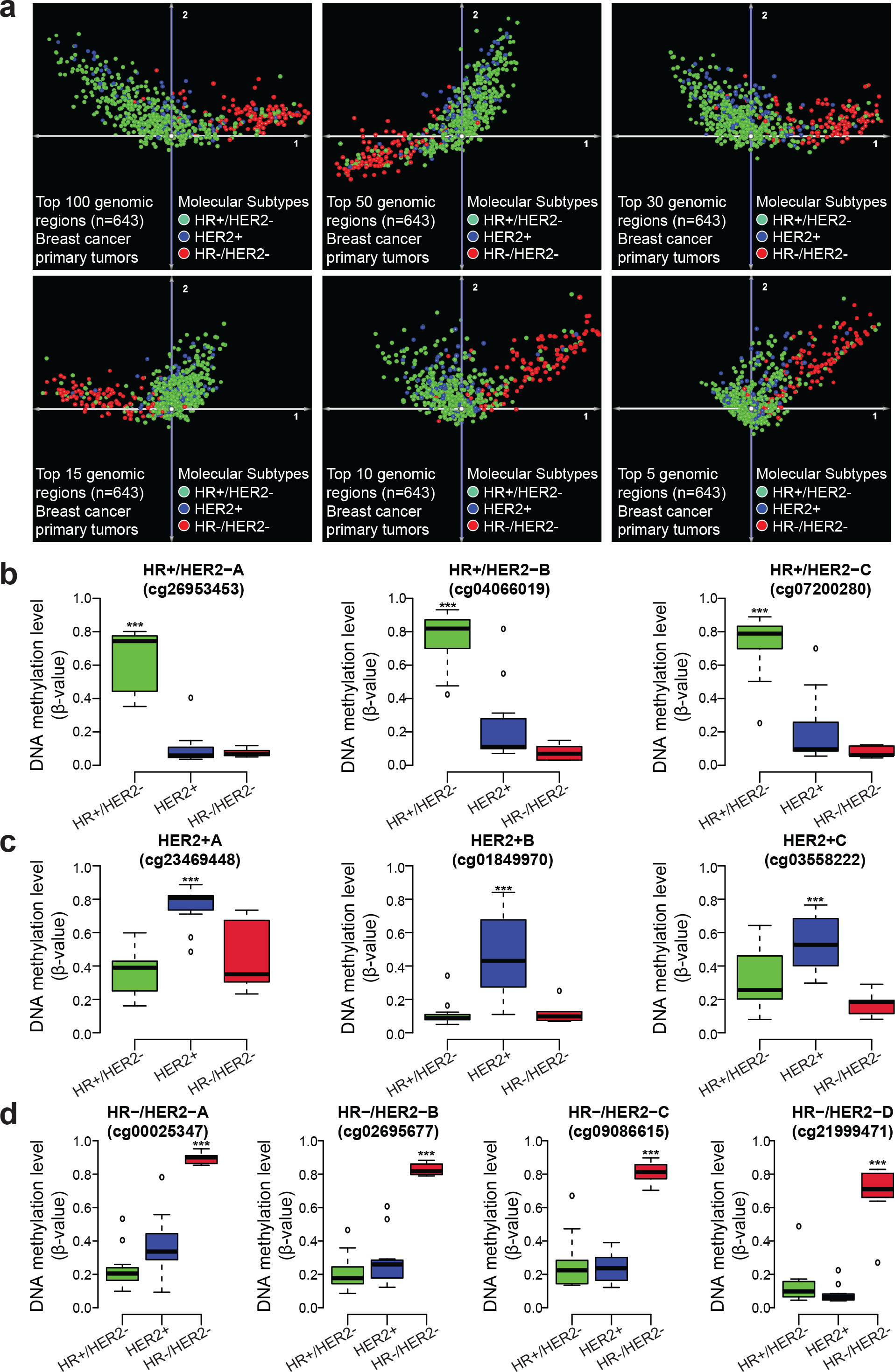
**a-** Principal component analyses for primary breast cancer using the top 100, 50, 30, 15, 10, and 5 most informative genomic regions for the classification of breast cancer subtypes. Only TCGA primary breast tumor specimens detection P-value greater than 0.01 for the 100 selected CpG sites were included in this analysis (n=643). **b-d** Boxplots showing the DNAm levels (HM450K microarray β-values) of 10 genomic regions differentially methylated among the three breast cancer brain metastases molecular subtypes (see **Supplementary Table 8** for details about the genomic location and distance to nearby genes). This set of regions includes three CpG sites hypermethylated in HR+ and HER2- breast cancer brain metastasis specimens (**b**), three CpG sites hypermethylated in HER2+ breast cancer brain metastasis specimens (**c**), and four CpG sites hypermethylated in HR- and HER2- breast cancer brain metastasis specimens (**d**). ***indicates a Wilcoxon’s test-based *P*-value less than or equal to 0.001.

**Supplementary Figure 7:**
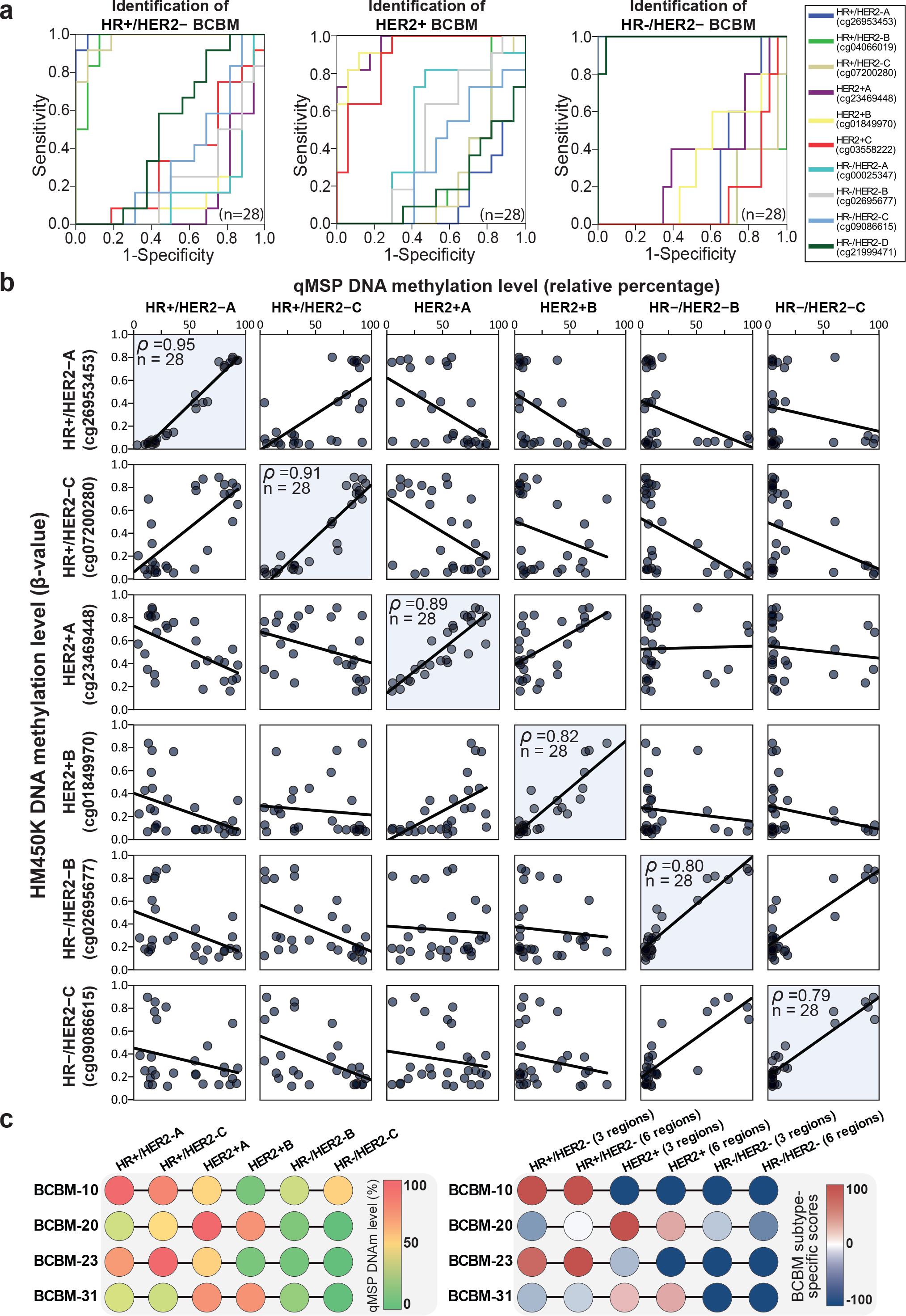
**a-** ROCs distinguishing among the three breast cancer brain metastasis molecular subtypes using the DNA methylation levels (HM450K microarray β-values) of the 10 differentially methylated regions (**Supplementary Table 8**). **b-** Spearman´s *ρ* correlation among DNA methylation levels assessed by HM450K (β-values, **y-axes**) and targeted approach (qMSP; **x-axes**) for the selected three genomic regions. **c-** qMSP analysis of the selected six genomic regions of the four BCBM samples without IHC assessment of ER, PgR, and HER2 at the moment of initial diagnosis. Left panel shows the DNAm level (percentage) for each region. Right panel shows the DNAm scores specific of each breast cancer molecular subtypes considering three (one per molecular subtype) or six (two per molecular subtypes) genomic regions. BCBM subtype-specific scores for the combination of three genomic regions were calculated as follow: **HR+/HER2-scores** = DNAm level of HR+/HER2- minus DNAm level of HER2+ minus DNAm level of HR-/HER2; **HER2+scores** = DNAm level of HER2+ minus DNAm level of HR+/HER2- minus DNAm level of HR-/HER2; and **HR-/HER2-scores** = DNAm level of HR-/HER2- minus DNAm level of HR+/HER2- minus DNAm level of HER2+. For the combination of six genomic regions, the average of regions specific of each BCBM subtype were considered.

**Supplementary Data 1:** Clinical and demographic information for all the patients with brain metastasis included in the study.

### Breast cancer brain metastasis (BCBM) specimens

#### **Specimen ID:** BCBM-01

Female patient with a diagnosis of invasive breast cancer at 37 years old. She developed a single cerebellar brain metastasis followed by lung and liver metastases. The immunohistochemistry markers of the brain metastasis were ER negative, PR negative and HER2 positive.

#### **Specimen ID:** BCBM-02

Female patient with a diagnosis of occult breast cancer with liver metastasis at 50 years old. Four years later, she developed multiple cerebellar and frontal brain metastases. The immunohistochemistry markers of the brain metastases were ER negative, PR negative and HER2 positive.

#### Specimens ID: BCBM-03 and BCBM-04

Female patient with a diagnosis of triple negative breast cancer at 34 years old. She initially developed lung and bone metastases, followed by 2 synchronous brain metastases in the frontal and parietal lobes, at 52 years old. The immunohistochemistry markers of the frontal lesion were ER negative, PR negative and HER2 negative.

#### Specimens ID: BCBM-05 and BCBM-19

Female patient with a diagnosis of invasive breast cancer at 50 years old. Two years later, she developed a frontal brain metastasis, followed by a subsequent parietal brain metastasis 3 months later. The immunohistochemistry markers of the first brain metastasis were ER negative, PR negative and HER2 positive.

#### **Specimen ID:** BCBM-06

Female patient with a diagnosis of invasive breast cancer at 47 years old. She presented bone and liver metastases, followed by lung metastases. She finally developed a single parietal-occipital brain metastasis. The immunohistochemistry markers of the brain metastasis were ER positive, PR positive and HER2 positive.

#### **Specimen ID:** BCBM-07

Female patient with a diagnosis of invasive ductal breast cancer, stage IA, at 76 years old. She presented a single cerebellar metastasis 4 years after the diagnosis of the primary breast cancer. The immunohistochemistry markers of the brain metastasis were ER negative, PR negative and HER2 negative.

#### **Specimen ID:** BCBM-08

Female patient with a diagnosis of invasive ductal breast cancer, stage IA, at 40 years old. She initially developed lung and bone metastases, followed by a single parietal brain metastasis at 50 years old. The immunohistochemistry markers of the brain metastasis were ER positive, PR positive and HER2 positive.

#### **Specimen ID:** BCBM-09

Female patient with a diagnosis of triple negative breast cancer, at 24 years old. She initially developed poorly differentiated metastasis at the chest wall, mediastinum lymph nodes and bone, followed by multiple brain metastases. The immunohistochemistry markers of the brain metastases were ER negative, PR negative and HER2 negative.

#### **Specimen ID:** BCBM-10

Female patient with a diagnosis of occult breast cancer with bone metastasis at 75 years old. Two years later, she developed a single brain metastasis at temporal lobe. The immunohistochemistry markers of the brain metastasis were retrospectively examined during this study and were ER positive, PR positive and HER2 negative, in concordance with the immunohistochemistry profile of the previous bone metastasis.

#### **Specimen ID:** BCBM-11

Female patient with a diagnosis of invasive ductal carcinoma triple negative breast cancer, stage IIA, at 38 years old. She initially developed lung and liver metastases, followed by multiple brain metastases at 40 years old. The immunohistochemistry markers of the brain metastases were ER negative, PR negative and HER2 negative.

#### **Specimen ID:** BCBM-12

Female patient with a diagnosis of invasive breast cancer, at 48 years old. She initially developed metastasis in the mediastinum, followed by multiple brain metastases at 50 years old. The immunohistochemistry markers of the brain metastases were ER positive, PR negative and HER2 positive.

#### **Specimen ID:** BCBM-13

Female patient with a diagnosis of inflammatory breast cancer, stage IIIB, at 50 years old. She developed a single brain metastasis in cerebellum 6 years after the diagnosis of the primary breast cancer. The immunohistochemistry markers of the brain metastasis were ER negative, PR negative and HER2 negative.

#### **Specimen ID:** BCBM-14

Female patient with a diagnosis of invasive breast cancer, at 39 years old. She initially developed bone, lung and liver metastases, followed by multiple brain metastases at 44 years old. The immunohistochemistry markers of the brain metastases were ER positive, PR negative and HER2 negative.

#### **Specimen ID:** BCBM-15

Female patient with a diagnosis of mucinous breast cancer, stage IIA, at 60 years old. She developed bone metastases followed by a single brain metastasis in temporal lobe 5 years after the diagnosis of the primary breast cancer. The immunohistochemistry markers of the brain metastasis were ER positive, PR positive and HER2 negative.

#### **Specimen ID:** BCBM-16

Female patient with a diagnosis of occult breast cancer with liver and cerebellum metastases at 73 years old. The immunohistochemistry markers of the brain metastases were ER positive, PR positive and HER2 negative.

#### **Specimen ID:** BCBM-17

Female patient with a diagnosis of invasive breast cancer, stage IIIC, at 37 years old. She developed parietal brain metastases 3 years after the diagnosis of the primary breast cancer. The immunohistochemistry markers of the brain metastases were ER positive, PR negative and HER2 negative.

#### **Specimen ID:** BCBM-20

Female patient with a diagnosis of *de novo* stage IV breast cancer, due to the presence of bone, lung and liver metastases, at 55 years old. Two years later, she developed a single cerebellum metastasis. The immunohistochemistry markers were retrospectively examined during this study and were ER negative, PR negative and HER2 positive.

#### **Specimen ID:** BCBM-21

Female patient with a diagnosis of invasive lobular breast cancer, stage IIA, at 43 years old. She developed frontal and temporal brain metastases 3 years after the diagnosis of the primary breast cancer. The immunohistochemistry markers of the brain metastases were ER positive, PR positive and HER2 negative.

#### **Specimen ID:** BCBM-22

Female patient with a diagnosis of invasive breast cancer, stage IIB, at 58 years old. She developed lymph node, bone and liver metastases and subsequently presented multiple brain metastases at 70 years old. The immunohistochemistry markers of the brain metastases were ER positive, PR negative and HER2 negative.

#### **Specimen ID:** BCBM-23

Female patient with a diagnosis of invasive ductal breast cancer, stage IIIA, at 36 years old. Five years later, she developed metastatic disease at mediastinum, bone, and lung. At the age of 46 years, she presented multiple cerebellum metastases. The immunohistochemistry markers were retrospectively examined during this study and were ER positive, PR negative and HER2 negative, similar to the immunohistochemistry profile of the primary breast cancer.

#### **Specimen ID:** BCBM-24

Female patient with a diagnosis of invasive ductal breast cancer, stage IB, at 59 years old. Two years later, she presented a chest wall recurrence. At the age of 71 years, she presented lung metastases followed by multiple brain metastases at frontal and occipital lobes. The immunohistochemistry markers of the brain metastases were ER positive, PR positive and HER2 positive.

#### **Specimen ID:** BCBM-25

Female patient with a diagnosis of occult breast cancer with bone and lung metastases at 55 years old. She developed multiple brain metastases at 67 years old. The immunohistochemistry markers of the brain metastases were ER positive, PR positive and HER2 negative.

#### **Specimen ID:** BCBM-26

Female patient with a diagnosis of invasive breast cancer, at 54 years old. Subsequently, she developed a single brain metastasis in the thalamus 4 years after the diagnosis of the primary breast cancer. The immunohistochemistry markers of the brain metastasis were ER negative, PR negative and HER2 negative.

#### **Specimen ID:** BCBM-27

Female patient with a diagnosis of brain metastasis due to breast cancer, at 61 years old. The immunohistochemistry markers of the brain metastasis were ER positive, PR positive and HER2 negative.

#### **Specimen ID:** BCBM-28

Female patient with a diagnosis of brain metastasis due to breast cancer, at 65 years old. The immunohistochemistry markers of the brain metastasis were ER positive, PR positive and HER2 negative.

#### **Specimen ID:** BCBM-31

Female patient with a diagnosis of invasive breast cancer, at 33 years old. Two years later, she developed a brain metastasis. The immunohistochemistry markers were retrospectively examined during this study and were ER positive, PR positive and HER2 positive.

#### **Specimen ID:** BCBM-32

Female patient with a diagnosis of brain metastasis due to breast cancer, at 45 years old. The immunohistochemistry markers of the brain metastasis were ER positive, PR positive and HER2 positive.

#### **Specimen ID:** BCBM-33

Female patient with a diagnosis of invasive ductal breast cancer, stage IIB, at 38 years old. Two years later, she presented cerebellum metastases. The immunohistochemistry markers of the brain metastases were ER negative, PR negative and HER2 positive.

### Lung cancer brain metastasis (LCBM) specimens

#### **Specimen ID:** LCBM-01

Female patient with a diagnosis of non-small cell lung cancer at 59 years old. She developed brain metastasis at 62 years old, similar to the previous bronchial biopsy.

#### **Specimen ID:** LCBM-02

Female patient with a diagnosis of non-small cell lung cancer at 63 years old. She developed brain metastasis at 65 years old, similar to the previous right lower lobe lung tumor. Immunohistochemistry markers were TTF-1 positive, Napsin-A positive.

#### **Specimen ID:** LCBM-03

Female patient with a diagnosis of non-small cell lung cancer with adenosquamous histology at 74 years old. Six months later, she developed brain metastasis similar to lung cancer with adenosquamous histology. Immunohistochemistry markers were TTF-1 positive, Napsin-A positive.

#### **Specimen ID:** LCBM-04

Female patient with a history of non-small cell lung cancer. She developed brain metastasis at 71 years old, compatible with pulmonary adenocarcinoma metastasis with neuroendocrine differentiation.

#### **Specimen ID:** LCBM-06

Female patient with a history of non-small cell lung cancer. She developed brain metastasis at 67 years old, compatible with metastatic adenocarcinoma of lung/bronchogenic origin. The brain metastasis was KRAS and EGFR wild-type.

#### **Specimen ID:** LCBM-08

Female patient with a history of non-small cell lung cancer at 66 years old. She developed brain metastasis 5 months later, with identical immunoprofile to primary lung carcinoma.

#### **Specimen ID:** LCBM-09

Female patient with a history of undifferentiated non-small cell lung cancer, presenting brain metastasis of similar histology at 66 years old.

#### **Specimen ID:** LCBM-10

Female patient with a diagnosis of brain metastasis from small cell lung cancer, at 71 years old. Immunohistochemistry markers were TTF-1 negative, synaptophysin positive, CK7 positive/CK20 negative.

#### **Specimen ID:** LCBM-12

Female patient with a diagnosis of non-small cell lung cancer with squamous histology at 86 years old. She developed brain metastasis 7 months later, with identical immunoprofile to primary lung carcinoma.

#### **Specimen ID:** LCBM-14

Male patient with a diagnosis of brain metastasis from small cell lung cancer, at 67 years old. Immunohistochemistry markers were TTF-1 positive, synaptophysin positive and chromogranin positive.

#### **Specimen ID:** LCBM-15

Male patient with a diagnosis of brain metastasis from non-small cell lung cancer with adenocarcinoma histology, at 88 years old. Immunohistochemistry markers were TTF-1 positive, napsin-A positive, and CK7 positive/CK20 negative.

#### **Specimen ID:** LCBM-16

Female patient with a diagnosis of brain metastasis from non-small cell lung cancer with adenocarcinoma histology, at 69 years old. Immunohistochemistry markers were TTF-1 positive, napsin-A positive, and CK7 positive/CK20 negative.

#### **Specimen ID:** LCBM-17

Female patient with a diagnosis of brain metastasis from non-small cell lung cancer with adenocarcinoma histology, at 64 years old.

#### **Specimen ID:** LCBM-18

Male patient with a diagnosis of brain metastasis of unknown primary at 61 years old. He has a history of skin basal cell carcinoma and melanoma in situ. The immunohistochemistry markers were compatible with pulmonary metastatic adenocarcinoma.

#### **Specimen ID:** LCBM-19

Female patient with a diagnosis of non-small cell lung cancer with adenocarcinoma histology at 81 years old. She developed brain metastasis 2 years later, with similar immunoprofile to primary lung carcinoma.

#### **Specimen ID:** LCBM-20

43 years old male patient with a history of an apical lung mass, presenting a single brain metastasis with histology of high-grade neuroendocrine carcinoma, synaptophysin positive, chromogranin positive and CK7 positive.

#### **Specimen ID:** LCBM-21

76 years old female patient with a diagnosis of brain metastasis compatible with lung adenocarcinoma origin. Immunohistochemistry markers were TTF-1 positive, napsin-A positive, and CK7 positive/CK20 negative. She has a history of a lung mass, heavier smoker. The lung biopsy couldn’t be performed due to patient’s poor health condition.

#### **Specimen ID:** LCBM-22

Female patient with a diagnosis of brain metastasis from non-small cell lung cancer with adenocarcinoma histology, at 68 years old.

### Melanoma brain metastasis (MBM) specimens

#### **Specimen ID:** MBM-09

Male patient diagnosed with stage IIB melanoma on the scalp at 60 years old. He developed scalp recurrence and regional lymph node metastasis, followed by lung metastasis. Finally, presented a single sellar metastasis, NRAS mutated, 73 months after the diagnosis of the primary tumor. The patient was deceased 25 months after the brain metastasis diagnosis.

#### **Specimen ID:** MBM-10

Female patient diagnosed with stage IIIB melanoma in the trunk at 51 years old. She developed cutaneous metastasis, and finally, presented a single brain metastasis at right basal ganglia, BRAF and NRAS wild-type, 12 years after the diagnosis of the primary tumor. The patient was deceased 43 months after the brain metastasis diagnosis.

#### **Specimen ID:** MBM-11

Male patient diagnosed with occult melanoma at 80 years old with simultaneous lung and brain metastasis at initial diagnosis. He presented multiple brain metastases at frontal, temporal lobes and cerebellum. The patient was deceased 6 months after the brain metastasis diagnosis.

#### **Specimen ID:** MBM-12

Male patient diagnosed with occult melanoma at 43 years old, presenting 2 brain metastases at left parietal and left temporal-parietal lobes, NRAS mutated. The patient was deceased 24 months after the brain metastasis diagnosis.

#### **Specimen ID:** MBM-14

Male patient, with a diagnosis of nodular melanoma on the scalp stage IIIC at 60 years-old. He subsequently developed cutaneous metastasis, and finally, presented a single brain metastasis at the left frontal lobe, BRAF mutated, 20 months after the diagnosis of the primary tumor. The patient was deceased 11 months after the brain metastasis diagnosis.

#### **Specimen ID:** MBM-15

Female patient, with a diagnosis of melanoma on the trunk at 44 years-old who presented 2 brain metastases at the left frontal lobe, NRAS mutated, 11 months after the diagnosis of the primary tumor. The patient is alive with persistent disease 5.6 years after the brain metastasis diagnosis.

#### **Specimen ID:** MBM-16

Female patient, with a diagnosis of melanoma at 44 years-old who presented a single brain metastasis at the right frontal lobe, BRAF mutated, 4 months after the initial diagnosis. The patient is alive with persistent disease 3 years after the brain metastasis diagnosis.

#### **Specimen ID:** MBM-17

Female patient, with a diagnosis of superficial spreading melanoma on the trunk who presented a single brain metastasis at the left parietal lobe, BRAF mutated, 21.6 years after the initial diagnosis. The patient was deceased 11 months after the brain metastasis diagnosis.

#### **Specimen ID:** MBM-18

75 years old Female patient, diagnosed with stage IIB nodular melanoma in the calf. She presented a single brain metastasis at the left frontal lobe, NRAS mutated, 10 years after primary diagnosis. The patient was deceased 5 months after the brain metastasis diagnosis.

#### **Specimen ID:** MBM-19

Male patient, diagnosed with melanoma on the face, stage IB at 54 years-old. He subsequently developed cutaneous and regional lymph nodes metastasis, followed by liver metastasis. Finally, presented 2 brain metastases at left Sylvian fissure, BRAF mutated, 22 months after the diagnosis of the primary tumor. The patient died 17 months after the brain metastasis diagnosis.

#### **Specimen ID:** MBM-20

Female patient, with a diagnosis of desmoplastic melanoma on the upper arm stage IIB at 62 years-old. She developed lung metastasis and 8 years and 10 months after the diagnosis of the primary melanoma. She presented a single brain metastasis at the right frontal lobe, BRAF/NRAS wild type. The patient is alive with persistent disease 16 months after the brain metastasis diagnosis.

#### **Specimen ID:** MBM-21

Female patient, with a diagnosis of melanoma at 56 years-old, stage IIIC. She presented a single brain metastasis at the left parietal lobe, BRAF/NRAS wild-type, 30 months after the diagnosis of the primary tumor. The patient died 3 years after the diagnosis of brain metastasis.

#### **Specimen ID:** MBM-22

Male patient, with a diagnosis of nodular melanoma on the face stage IIA at 48 years old. He presented a single brain metastasis at the right temporal region, NRAS mutated, 52 months after the diagnosis of the primary tumor. He previously developed bone and lung metastases. The patient died 3 years after the brain metastasis diagnosis from a non-caner related death.

#### **Specimen ID:** MBM-23

Male patient, with a diagnosis of melanoma on the shoulder, stage IIB, at 70 years old. He subsequently developed right posterior chest cutaneous metastasis, and finally, presented a single brain metastasis at the right frontal lobe, NRAS mutated, 28 months after the diagnosis of the primary tumor. The patient was deceased 11 months after the brain metastasis diagnosis.

#### **Specimen ID:** MBM-24

Male patient, with a diagnosis of melanoma at 65 years-old, stage II. He subsequently developed liver, lung and bone metastases, and finally, presented a single brain metastasis at posterior left inferior gyrus lesion, BRAF/NRAS wild-type, 6 years after the diagnosis of the primary tumor. The patient was deceased 4 months after the brain metastasis diagnosis.

#### **Specimen ID:** MBM-25

Male patient, with a diagnosis of stage IIA thoracic melanoma at 60 years-old who subsequently developed 2 brain metastases at right upper parietal, BRAF mutated, 23 months after the diagnosis of the primary tumor. The patient was deceased 7.8 years after the brain metastasis diagnosis.

#### **Specimen ID:** MBM-26

Male patient, with a diagnosis of melanoma on the flank stage IIA at 69 years-old. He then developed lung metastasis, and finally, presented a single brain metastasis at right temporal, NRAS mutated, 5 years after the diagnosis of the primary tumor. The patient was deceased 7 months after the brain metastasis diagnosis.

#### **Specimen ID:** MBM-27

71-year-old male patient diagnosed with stage IIA melanoma on the shoulder. He subsequently developed bone metastasis, and finally, presented a single brain metastasis at the left temporal parietal region, BRAF mutated, 11 years after the diagnosis of the primary tumor. The patient was deceased 6 months after the brain metastasis diagnosis.

#### **Specimen ID:** MBM-28

Male patient, with a thoracic nodular melanoma stage IIC at 44 years-old. He presented a single brain metastasis at right parietal-occipital, BRAF mutated, 31 months after the diagnosis of the primary tumor. The patient is alive and disease-free 8 years after the brain metastasis diagnosis.

#### **Specimen ID:** MBM-29

Male patient, diagnosed with stage IIA melanoma on the forehead, at 74 years-old. He presented a single brain metastasis at the right occipital lobe, NRAS mutated. The patient was deceased 9 months after the brain metastasis diagnosis.

#### **Specimen ID:** MBM-30

Male patient diagnosed with stage IIB thoracic desmoplastic melanoma at 61 years-old. He presented 2 brain metastases in the right occipital lobe, BRAF/NRAS wild-type, 38 months after the diagnosis of the primary tumor. The patient is alive with persistent disease 5 months after the brain metastasis diagnosis.

#### **Specimen ID:** MBM-31

Male patient diagnosed with stage IIIC melanoma on the trunk at 65 years-old. He subsequently developed lung and regional lymph node metastasis, and finally, presented 2 brain metastases at the left frontal lobe, NRAS mutated, 6 years after the diagnosis of the primary tumor. The patient was deceased 3 months after the brain metastasis diagnosis.

#### **Specimen ID:** MBM-32

42-year-old female patient, with a diagnosis of stage IB superficial spreading melanoma on the trunk. She subsequently developed right neck lymph node metastasis, and finally, presented a single brain metastasis at the left temporal lobe, BRAF mutated, 58 months after the diagnosis of the primary tumor. The patient was deceased 3.9 years after the brain metastasis diagnosis.

#### **Specimen ID:** MBM-33

Male patient, with a diagnosis of stage IIB nodular melanoma on the trunk at 60 years-old. He subsequently developed cutaneous metastasis followed by liver metastasis. Finally, presenting a single brain metastasis at the left frontal lobe, BRAF mutated, 20 months after the diagnosis of the primary tumor. The patient was deceased 11 months after the brain metastasis diagnosis.

#### **Specimen ID:** MBM-34

Male patient diagnosed with stage IIA superficial spreading melanoma on the head/neck at 52 years old, who presented two brain metastases at right parietal-occipital, NRAS mutated 34 months after primary diagnosis. The patient was deceased 3 months after brain metastases diagnosis.

#### **Specimen ID:** MBM-35

Male patient, with a diagnosis of stage IIB nodular melanoma on the scalp at 61 years old. He subsequently developed cutaneous metastasis followed by lymph node metastasis. Finally presented a single brain metastasis at frontal cortex, BRAF mutated 17 months after the primary diagnosis. The patient was deceased 15 months after brain metastasis diagnosis.

#### **Specimen ID:** MBM-36

27-year-old male patient diagnosed with stage IB thoracic melanoma. He then developed right neck lymph node metastasis. Finally presented multiple brain metastases at right and left frontal cortex, BRAF mutated 12 years after the primary diagnosis. The patient is alive with persistent disease 6 years after brain metastases diagnosis.

#### **Specimen ID:** MBM-37

56-year-old male patient diagnosed with melanoma in situ on extremities. He subsequently presented right inguinal metastasis and finally developed a single brain metastasis in the left frontal lobe, NRAS mutated 2 years after the primary diagnosis. The patient died 22 months after brain metastasis diagnosis.

#### **Specimen ID:** MBM-38

Female patient with a diagnosis of stage IB superficial spreading melanoma on the shoulder at 38 years old. She subsequently developed right neck lymph node metastasis. Eleven years after the primary diagnosis she presented a single brain metastasis at the right frontal cortex, BRAF mutated. The patient is alive with persistent disease 7.2 years after the brain metastasis diagnosis.

#### **Specimen ID:** MBM-39

61-year-old male patient diagnosed with stage IIA superficial spreading melanoma on the extremities. He subsequently developed left inguinal lymph node metastasis, and finally presented a single brain metastasis at left occipital, BRAF mutated 3.5 years after the primary diagnosis. The patient was deceased 14 months after brain metastasis diagnosis.

#### **Specimen ID:** MBM-40

Male patient with a diagnosis of stage IA nodular melanoma on the trunk at 54 years old. He presented a single brain metastasis at left parietal, NRAS mutated 11 months after the initial diagnosis. The patient was deceased 11 months after brain metastasis diagnosis.

#### **Specimen ID:** MBM-41

Female patient with occult melanoma at 62 years old due to brain metastasis at right frontal-parietal, NRAS mutated. The patient was deceased 4 months after brain metastasis diagnosis.

#### **Specimen ID:** MBM-42

Male patient with stage IIIC melanoma on the foot at 17 years old. He then developed in transit right lower leg metastasis followed by bone metastasis. Finally presented two brain metastases at right cerebellar and right frontal lobe, BRAF mutated 14 years after initial diagnosis. The patient died 10 months after brain metastasis diagnosis.

#### **Specimen ID:** MBM-43

43-year-old male diagnosed with stage IB superficial spreading melanoma on the trunk who subsequently presented lung metastasis. He finally presented a single brain metastasis at frontal lobe, NRAS mutated 4.3 years after initial diagnosis. The patient was deceased 10 months after brain metastasis diagnosis.

#### **Specimen ID:** MBM-44

Male patient with a diagnosis of stage IA superficial spreading melanoma on the trunk at 68 years old. The patient subsequently developed regional lymph node metastasis followed by spleen metastasis. He finally developed a single brain metastasis at the left occipital lobe, BRAF mutated, 4 years after the initial diagnosis. The patient was deceased 12 months after brain metastasis diagnosis.

#### **Specimen ID:** MBM-45

40-year-old female patient diagnosed with stage IB lentigo maligna melanoma. She then presented multiple simultaneous brain, right chest, small intestine, and multiple lymphadenopathies metastases. The multiple brain metastases were located in right frontal, and cerebellum, BRAF mutated.

#### **Specimen ID:** MBM-46

75-year-old male patient diagnosed with stage IA superficial spreading melanoma on mid-chest. He subsequently presented lumbar spine and retroperitoneal metastasis. He then developed four brain metastases at right frontal lobe and right temporoparietal, BRAF mutated 4.4 years after the initial diagnosis. The patient was deceased 3 months after brain metastases diagnosis.

#### **Specimen ID:** MBM-47

Male patient diagnosed with melanoma at 56 years old with a simultaneous brain, liver, and lung metastasis at initial diagnosis. He presented multiple brain metastases at right frontal-parietal, right occipital, right cerebellar, and left parietal, BRAF mutated. The patient died 11 months after brain metastasis diagnosis.

#### **Specimen ID:** MBM-48

Female patient diagnosed with melanoma at 50 years old. She subsequently developed shoulder, abdomen, chest, and back metastasis. 6 years after initial diagnosis the patient developed a single brain metastasis at left parietal, NRAS mutated. The patient was deceased 3 years after brain metastasis diagnosis.

#### **Specimen ID:** MBM-49

49-year-old male patient diagnosed with stage IB superficial spreading melanoma on extremities. He subsequently developed left knee, left groin lymph node followed by lung metastasis. Finally presenting two brain metastases at left hippocampus and left frontal lobe, NRAS mutated 3 years after initial diagnosis. The patient died 2 years after brain metastasis diagnosis.

#### **Specimen ID:** MBM-50

Male patient diagnosed with melanoma on the trunk at age 41. He developed cutaneous, lung, liver, and abdomen metastasis with lymph node metastasis. He then developed a single brain metastasis at right temporal, BRAF mutated 3 years after initial diagnosis. The patient died 16 months after brain metastasis diagnosis.

#### **Specimen ID:** MBM-51

Male patient diagnosed with occult melanoma at 76 years old. He developed pulmonary hillium followed by liver metastasis. 15 months after the initial diagnosis he presented a single brain metastasis at the left frontal lobe. The patient died 1 month after brain metastasis diagnosis.

#### **Specimen ID:** MBM-52

Male patient diagnosed with occult melanoma due to lung metastasis, at 71 years old. He then presented a single brain metastasis at right occipital, d-WT 19 months after initial diagnosis. The patient died 11 months after brain metastasis diagnosis.

#### **Specimen ID:** MBM-53

51-year-old male patient diagnosed with stage IIB melanoma with simultaneous brain and lung metastasis followed by duodenal adenocarcinoma metastasis at initial diagnosis. He then presented a single brain metastasis at right parietal, NRAS mutated 3 years after the initial diagnosis. The patient died 7 years after brain metastasis diagnosis.

### Brain metastasis (BM) specimens with unknown or uncertain diagnosis

#### **Specimen ID:** BM-01

Female patient, with a history of breast and lung cancer at 54 years old. She developed brain metastases at 58 years old. Anatomic-pathology evaluation compatible with lung cancer brain metastasis. No confirmatory immunohistochemistry evaluation was performed.

#### **Specimen ID:** BM-02

Female patient, with a history of invasive breast cancer, but no history of lung cancer. She developed brain metastases at 68 years old. Immunohistochemistry evaluation compatible with lung cancer brain metastasis (TTF-1 positive, Napsin-A positive).

#### **Specimen ID:** BM-03

Female patient with a history of non-small cell lung cancer at 43 years, invasive breast cancer and, non-Hodgkin’s lymphoma at 71 years old. At 80 years old, she presented a brain metastasis in the frontal lobe, compatible with metastatic pulmonary adenocarcinoma.

#### **Specimen ID:** BM-04

64 years old female patient with a history of pulmonary adenocarcinoma. She developed brain metastases histologically compatible with metastatic pulmonary adenocarcinoma, but lacking confirmatory immunohistochemistry, CK7positive/CK20 negative, TTF-1 negative, Napsin-A negative.

## References

1 Schouten, L. J., Rutten, J., Huveneers, H. A. & Twijnstra, A. Incidence of brain metastases in a cohort of patients with carcinoma of the breast, colon, kidney, and lung and melanoma. Cancer 94, 2698–2705 (2002).

2 Barnholtz-Sloan, J. S. et al. Incidence proportions of brain metastases in patients diagnosed (1973 to 2001) in the Metropolitan Detroit Cancer Surveillance System. J. Clin. Oncol. 22, 2865–2872 (2004).

3 Berghoff, A. S. et al. Descriptive statistical analysis of a real life cohort of 2419 patients with brain metastases of solid cancers. ESMO Open 1, e000024 (2016).

4 Cagney, D. N. et al. Incidence and prognosis of patients with brain metastases at diagnosis of systemic malignancy: A population-based study. Neuro Oncol. (2017).

5 Gavrilovic, I. T. & Posner, J. B. Brain metastases: epidemiology and pathophysiology. J. Neurooncol. 75, 5–14 (2005).

6 Lin, X. & DeAngelis, L. M. Treatment of Brain Metastases. J. Clin. Oncol. 33, 3475–3484 (2015).

7 Ramakrishna, N. et al. Recommendations on disease management for patients with advanced human epidermal growth factor receptor 2-positive breast cancer and brain metastases: American Society of Clinical Oncology clinical practice guideline. J. Clin. Oncol. 32, 2100–2108 (2014).

8 Costa, R. et al. Developmental therapeutics for patients with breast cancer and central nervous system metastasis: current landscape and future perspectives. Ann. Oncol. 28, 44–56 (2017).

9 Bekaert, L., Emery, E., Levallet, G. & Lechapt-Zalcman, E. Histopathologic diagnosis of brain metastases: current trends in management and future considerations. Brain Tumor Pathol. 34, 8–19 (2017).

10 Becher, M. W., Abel, T. W., Thompson, R. C., Weaver, K. D. & Davis, L. E. Immunohistochemical analysis of metastatic neoplasms of the central nervous system. J. Neuropathol. Exp. Neurol. 65, 935–944 (2006).

11 Hyman, D. M., Taylor, B. S. & Baselga, J. Implementing Genome-Driven Oncology. Cell 168, 584–599 (2017).

12 Moran, S. et al. Epigenetic profiling to classify cancer of unknown primary: a multicentre, retrospective analysis. Lancet Oncol. 17, 1386–1395 (2016).

13 Marzese, D. M. et al. DNA methylation and gene deletion analysis of brain metastases in melanoma patients identifies mutually exclusive molecular alterations. Neuro Oncol. 16, 1499–1509 (2014).

14 Marzese, D. M. et al. Epigenome-wide DNA methylation landscape of melanoma progression to brain metastasis reveals aberrations on homeobox D cluster associated with prognosis. Hum. Mol. Genet. 23, 226–238 (2014).

15 Marzese, D. M., Huynh, J. L., Kawas, N. P. & Hoon, D. S. Multi-platform Genome-wide Analysis of Melanoma Progression to Brain Metastasis. Genom. Data 2, 150–152 (2014).

16 Marzese, D. M. et al. Brain metastasis is predetermined in early stages of cutaneous melanoma by CD44v6 expression through epigenetic regulation of the spliceosome. Pigment Cell Melanoma Res. 28, 82–93 (2015).

17 The Cancer Genome Atlas Network. Comprehensive molecular portraits of human breast tumours. Nature 490, 61–70 (2012).

18 Genomic Classification of Cutaneous Melanoma. Cell 161, 1681–1696 (2015).

19 Zhou, W., Laird, P. W. & Shen, H. Comprehensive characterization, annotation and innovative use of Infinium DNA methylation BeadChip probes. Nucleic Acids Res. 45, e22–e22 (2017).

20 Sandoval, J. et al. Validation of a DNA methylation microarray for 450,000 CpG sites in the human genome. Epigenetics 6, 692–702 (2011).

21 Chen, Y. A. et al. Discovery of cross-reactive probes and polymorphic CpGs in the Illumina Infinium HumanMethylation450 microarray. Epigenetics 8, 203–209 (2013).

22 Moran, S., Arribas, C. & Esteller, M. Validation of a DNA methylation microarray for 850,000 CpG sites of the human genome enriched in enhancer sequences. Epigenomics 8, 389–399 (2016).

23 Bauer, A. H., Erly, W., Moser, F. G., Maya, M. & Nael, K. Differentiation of solitary brain metastasis from glioblastoma multiforme: a predictive multiparametric approach using combined MR diffusion and perfusion. Neuroradiology 57, 697–703 (2015).

24 Brennan, C. W. et al. The somatic genomic landscape of glioblastoma. Cell 155, 462–477 (2013).

25 Aran, D., Sirota, M. & Butte, A. J. Systematic pan-cancer analysis of tumour purity. Nat. Commun. 6, 8971 (2015).

26 McLean, C. Y. et al. GREAT improves functional interpretation of cis-regulatory regions. Nat. Biotechnol. 28, 495–501 (2010).

27 Badve, S. et al. Basal-like and triple-negative breast cancers: a critical review with an emphasis on the implications for pathologists and oncologists. Mod. Pathol. 24, 157–167 (2011).

28 Prat, A. & Perou, C. M. Mammary development meets cancer genomics. Nat. Med. 15, 842–844 (2009).

29 Hoadley, K. A. et al. Multiplatform analysis of 12 cancer types reveals molecular classification within and across tissues of origin. Cell 158, 929–944 (2014).

30 Breiman, L. Random forests. Machine learning 45, 5–32 (2001).

31 Pekmezci, M. & Perry, A. Neuropathology of brain metastases. Surg. Neurol. Int. 4, S245–255 (2013).

32 Takei, H., Rouah, E. & Ishida, Y. Brain metastasis: clinical characteristics, pathological findings and molecular subtyping for therapeutic implications. Brain Tumor Pathol. 33, 1–12 (2016).

33 Euskirchen, P. et al. Same-day genomic and epigenomic diagnosis of brain tumors using real-time nanopore sequencing. Acta Neuropathol. 134, 691–703 (2017).

34 Chamberlain, M. C., Baik, C. S., Gadi, V. K., Bhatia, S. & Chow, L. Q. Systemic therapy of brain metastases: non-small cell lung cancer, breast cancer, and melanoma. Neuro Oncol. 19, i1–i24 (2017).

35 Cardoso, F. et al. 3rd ESO-ESMO International Consensus Guidelines for Advanced Breast Cancer (ABC 3). Ann. Oncol. 28, 16–33 (2017).

36 Kaidar-Person, O. et al. Discrepancies between biomarkers of primary breast cancer and subsequent brain metastases: an international multicenter study. Breast Cancer Res. Treat. (2017).

37 Thomson, A. H. et al. Changing molecular profile of brain metastases compared with matched breast primary cancers and impact on clinical outcomes. Br. J. Cancer 114, 793–800 (2016).

38 Brastianos, P. K. et al. Genomic Characterization of Brain Metastases Reveals Branched Evolution and Potential Therapeutic Targets. Cancer Discov. 5, 1164–1177 (2015).

39 Hammond, M. E. et al. American Society of Clinical Oncology/College Of American Pathologists guideline recommendations for immunohistochemical testing of estrogen and progesterone receptors in breast cancer. J. Clin. Oncol. 28, 2784–2795 (2010).

40 Wolff, A. C. et al. Recommendations for human epidermal growth factor receptor 2 testing in breast cancer: American Society of Clinical Oncology/College of American Pathologists clinical practice guideline update. J. Clin. Oncol. 31, 3997–4013 (2013).

41 Maksimovic, J., Gordon, L. & Oshlack, A. SWAN: Subset-quantile Within Array Normalization for Illumina Infinium HumanMethylation450 BeadChips. Genome Biol. 13, R44 (2012).

42 Colaprico, A. et al. TCGAbiolinks: an R/Bioconductor package for integrative analysis of TCGA data. Nucleic Acids Res. 44, e71 (2016).

43 Saeed, A. I. et al. TM4 microarray software suite. Methods Enzymol. 411, 134–193 (2006).

